# Arginine methylation of DDX5 RGG/RG motif by PRMT5 regulates RNA:DNA resolution

**DOI:** 10.1101/451823

**Authors:** Sofiane Y. Mersaoui, Zhenbao Yu, Yan Coulombe, Martin Karam, Franciele F. Busatto, Jean-Yves Masson, Stéphane Richard

**Author notes:** Authors contributed equally (Sofiane Y. Mersaoui, Zhenbao Yu). Corresponding authors: Dr. Jean Yves Masson,; Dr. Stéphane Richard.

## Abstract

Aberrant transcription-associated RNA:DNA hybrid (R-loop) formation often lead to catastrophic conflicts during replication resulting in DNA double strand breaks and genome instability. To prevent such conflicts, these hybrids require dissolution by helicases and/or RNaseH. Little information is known about how these helicases are regulated. Herein, we identify DDX5, an RGG/RG motif containing DEAD-box family of RNA helicase, as a crucial player in R-loop resolution. We define at the mechanistic level the function of DDX5 in R-loop resolution. *In vitro*, recombinant DDX5 resolves R-loops in an ATP-dependent manner leading to R-loop degradation by the XRN2 exoribonuclease. DDX5 deficient cells accumulated R-loops at loci known to form R-loops using RNA:DNA immunoprecipitation (DRIP)-qPCR and increased RNaseH sensitive RAD51 foci. PRMT5, an arginine methyltransferase, associated with DDX5 and methylated its RGG/RG motif. This motif was required to associate with XRN2 and resolve cellular R-loops. Furthermore, PRMT5 deficient cells accumulated R-loops, as detected by DRIP-qPCR resulting in increased gH2AX foci. Our findings define a new mechanism by which an RNA helicase, DDX5, is modulated by arginine methylation to resolve R-loops.

## Introduction

In mammalian cells, there are nine protein arginine methyltransferases (PRMTs) responsible for transferring methyl groups from S-adenosyl-methionine to the nitrogen atoms of arginine (Bedford & Clarke, 2009). Arginine methylation modulates multiple biological processes from gene expression, pre-mRNA splicing, the DNA damage response and signal transduction (Blanc & Richard, 2017). PRMT5 is the major type enzyme catalyzing symmetrical dimethylarginines and as such multiple responses have been observed with inhibition of its function including the induction of the p53 response, DNA damage and cell death. PRMT5 regulates gene expression by methylating histones and transcription factors (Cho, Zheng et al., 2012, Friesen, Massenet et al., 2001, Pal, Vishwanath et al., 2004). Methylation of the Sm proteins regulates the p53 response by modulating the alternative splicing of *MDM4* and *MDM2* (Bezzi, Teo et al., 2013). Inhibition of arginine methylation has been shown to induce DNA damage and sensitize cells to DNA damaging agents, reviewed in (Blanc & Richard, 2017). PRMT5 regulates TIP60 activity by RUVBL1 methylation (Clarke, Sanchez-Bailon et al., 2017) and by TIP60 alternative splicing to regulate the DDR (Hamard, Santiago et al., 2018). This multitude of responses triggered in the absence of PRMT5 function suggest that it may be a good therapeutic target. Indeed PRMT5 inhibition, has been shown to decrease tumor growth in mouse models of mantle cell lymphoma, AML, CML, B cell lymphoma, glioma, and breast cancer (Braun, Stanciu et al., 2017, Chan-Penebre, Kuplast et al., 2015, Hamard et al., 2018, Kaushik, Liu et al., 2018, Koh, Bezzi et al., 2015, Li, Chitnis et al., 2015, Zhou, Xu et al., 2016). Herein, we show that the inhibition of PRMT5 accumulates R-loops associated with increased DNA damage.

Physiologically R-loops or RNA:DNA hybrids are programmed structures that occur during many cellular processes including transcription, replication, and immunoglobulin class switching (Bhatia, Herrera-Moyano et al., 2017, Chedin, 2016, Skourti-Stathaki & Proudfoot, 2014). Persistent R-loops impede DNA replication and if unresolved ultimately cause DNA breaks and genomic instability (Aguilera & Gomez-Gonzalez, 2017, Hamperl & Cimprich, 2016) or mitochondrial instability (Silva, Camino et al., 2018). Therefore, it is not surprising that there are specialized complexes to prevent and resolve R-loops. R-loops can be unwound by RNA:DNA helicases, such as Senataxin (SETX) and Aquarius (AQR) (Bhatia, Barroso et al., 2014, Hatchi, Skourti-Stathaki et al., 2015, Skourti-Stathaki, Proudfoot et al., 2011, Sollier, Stork et al., 2014) and the RNA in RNA:DNA hybrids degraded by RNaseH1 and RNaseH2 (Wahba, Amon et al., 2011). The RNA helicase Senataxin has been shown to resolve R-loops *in vitro* and its deficiency in cells leads to R-loop accumulation (Hatchi et al., 2015, Skourti-Stathaki et al., 2011, Sollier et al., 2014). Senataxin has been shown to function with the 5’-3’exonuclease XRN2 to resolve a subset of R-loops at transcription termination sites of actively transcribed genes (Aymard, Aguirrebengoa et al., 2017, Morales, Richard et al., 2016, Skourti-Stathaki et al., 2011). The DNA helicase RECQ5 and RNA helicases DDX1 (Li, Germain et al., 2016, Li, Monckton et al., 2008, Ribeiro de Almeida, Dhir et al., 2018), DDX19 (Hodroj, Recolin et al., 2017), DDX21 (Song, Hotz-Wagenblatt et al., 2017), DDX23 (Sridhara, Carvalho et al., 2017) and DHX9 (Cristini, Groh et al., 2018) were also found to be functionally involved in suppression of R-loops. Topoisomerase I removes the negative supercoils behind RNA polymerases to prevent annealing of the nascent RNA with the DNA template and suppresses R-loop formation (Tuduri, Crabbé et al., 2009). Fanconi anemia (FA) pathway proteins resolve RNA:DNA hybrids via FANCM translocase activity (García-Rubio, Pérez-Calero et al., 2015, Schwab, Nieminuszczy et al., 2015). Several RNA-processing proteins, such as the THO complex and the SRSF splicing factor, suppress R-loop formation namely by preventing the availability of the nascent RNAs for hybridization to template DNA (Huertas & Aguilera, 2003, Li & Manley, 2005, Paulsen, Soni et al., 2009, Sollier et al., 2014, Stirling, Chan et al., 2012, Wahba et al., 2011). The homologous recombination proteins, BRCA1 and BRCA2, are also involved in R-loop prevention and resolution (Bhatia et al., 2014, Hatchi et al., 2015). It is no surprise that mutations of proteins that prevent R-loop accumulation are frequently found in human diseases (Bhatia et al., 2017).

Little is known of the post-translational modifications regulating R-loop formation and resolution. It is known that the pausing of RNA polymerase II (RNA PolII) increases DDX23 phosphorylation by SRPK2 enhancing R-loop suppression (Sridhara et al., 2017). Acetylation of DDX21 by CBP regulates its helicase activity (Song et al., 2017). Methylation of RNA PolII subunit POLR2A by PRMT5 regulates Senataxin recruitment at transcription termination regions (Zhao, Gish et al., 2016), while methylation of TDRD3 by CARM1 regulates the recruitment of topoisomerase IIIB at c-Myc locus preventing negative supercoiling and R-loops (Yang, McBride et al., 2014). In this study, we define a new role for arginine methylation in the regulation of R-loops. We show that the methylation of the RNA helicase DDX5 by PRMT5 regulates its association with XRN2 to suppress R-loops and maintain genomic stability.

## Results

### DDX5 deficiency leads to an increase in RNaseH sensitive RAD51 foci.

A genome-wide screen identified RNA binding proteins including the DEAD box helicase DDX17 to localize to DNA damage sites (Adamson, Smogorzewska et al., 2012), however its function in the DNA double strand break (DSB) repair was not determined. DDX17 is often found in a complex with its ortholog DDX5 (Fuller-Pace, 2013). To investigate whether these RNA helicases play a role in the DNA damage response (DDR) pathway, we initially examined whether U2OS cells deficient in either helicase had altered ionizing radiation (IR)-induced DNA damage foci. Depletion of DDX5, but not DDX17, by siRNA transfection led to a significant increase of RAD51, RPA2 and BrdU foci at different time points after IR treatment (Figure 1A-C and S1A-D), while, DDX5 knockdown had little effect on γH2AX or 53BP1 foci formation (Figure S1E). In more details, we found that each RAD51 focus was much brighter (increasing from 1 to 4h) than in siCTL cells (Figure 1B and data not shown), suggesting that longer RAD51 filaments were forming next to each DSB. Moreover, there were more foci per cell at each time point, suggesting that without DDX5, RAD51 accumulated at lesions without completing repair. Next, we examined whether the helicase activity of DDX5 was required for RAD51 foci accumulation. DDX5 deficient cells were transfected with siRNA resistant expression vectors encoding FLAG epitope tagged wild type and helicase dead DDX5. Wild type DDX5, but not FLAG-DDX5 DEAD mutant, significantly restored RAD51 foci to lower levels, as observed in siCTL cells (Figure S2A, S2B), demonstrating that the DDX5 RNA helicase activity is required to maintain proper RAD51 foci dynamics.

**Figure S2.**
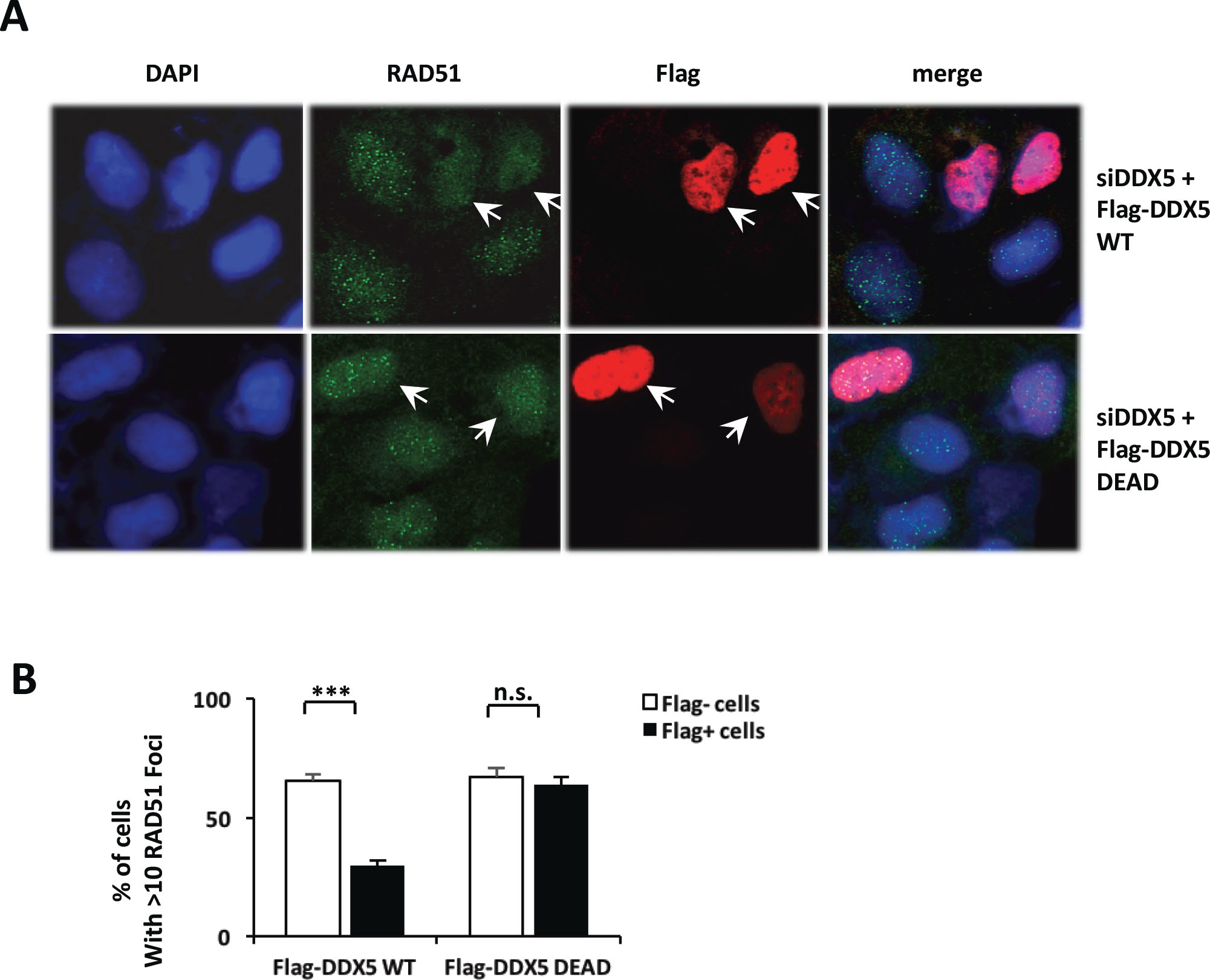
The DDX5 helicase activity is required to sustain DNA damage RAD51 foci. (A) U2OS cells were first transfected with siDDX5 siRNA (siDDX5 #1) and then the Flag-tagged DDX5 wild type or its catalytically inactive mutant (Flag-DDX5 DEAD) plasmid one day after the siRNA transfection. Two days after plasmid transfection, the cells were treated with 10 Gy of IR. After 4h of recovery, the cells were visualized by indirect immunofluorescence with anti-Flag (red) and anti-RAD51 (green) antibody. A typical image was shown for each sample. Arrows indicate the plasmid DNA-transfected cells. (B) The graph shows the average and standard error of the mean (S.E.M.) from three independent experiments performed in duplicates where >30 images were acquired containing at least 200 cells per condition. Statistical significance was assessed using Student’s t-test. *, *p* < 0.05; **, *p* < 0.01; ***, *p* < 0.001 and n.s., no significant.

**Figure S1.**
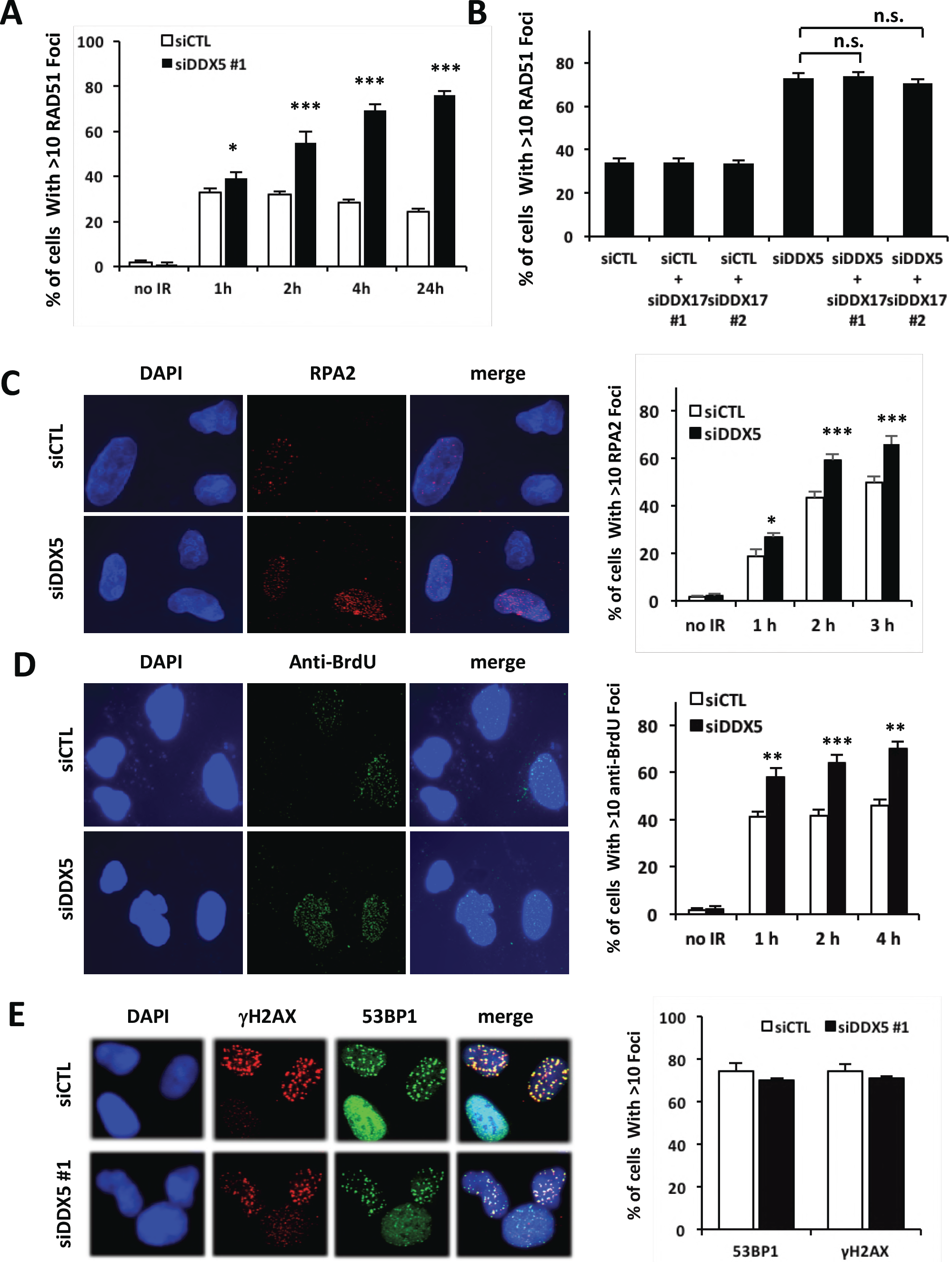
DDX5 siRNA transfected cells have sustained RAD51, RPA2 and BrdU foci after DNA damage, but not sustained gH2AX or 53BP1 foci. (A) U2OS cells were transfected with siDDX5 (#1) or control siRNA targeting luciferase (siCTL). The cells were left untreated or treated with 10 Gy of IR and allowed to recover for the indicated time or for 4 h (B). RAD51, RPA2 foci (C), anti-BrdU foci (D), gH2AX and 53BP1 foci (E) were analyzed as in the Materials and Methods. A typical image is shown for each sample. The cells with >10 foci were counted and expressed as a percentage. The graph shows the average and standard error of the mean (S.E.M.) from three independent experiments performed in duplicates where >30 images were acquired containing at least 200 cells per condition. Statistical significance was assessed using Student’s t-test. *, *p* < 0.05; **, *p* < 0.01; ***, *p* < 0.001 and n.s., no significant.

**Figure.1.**
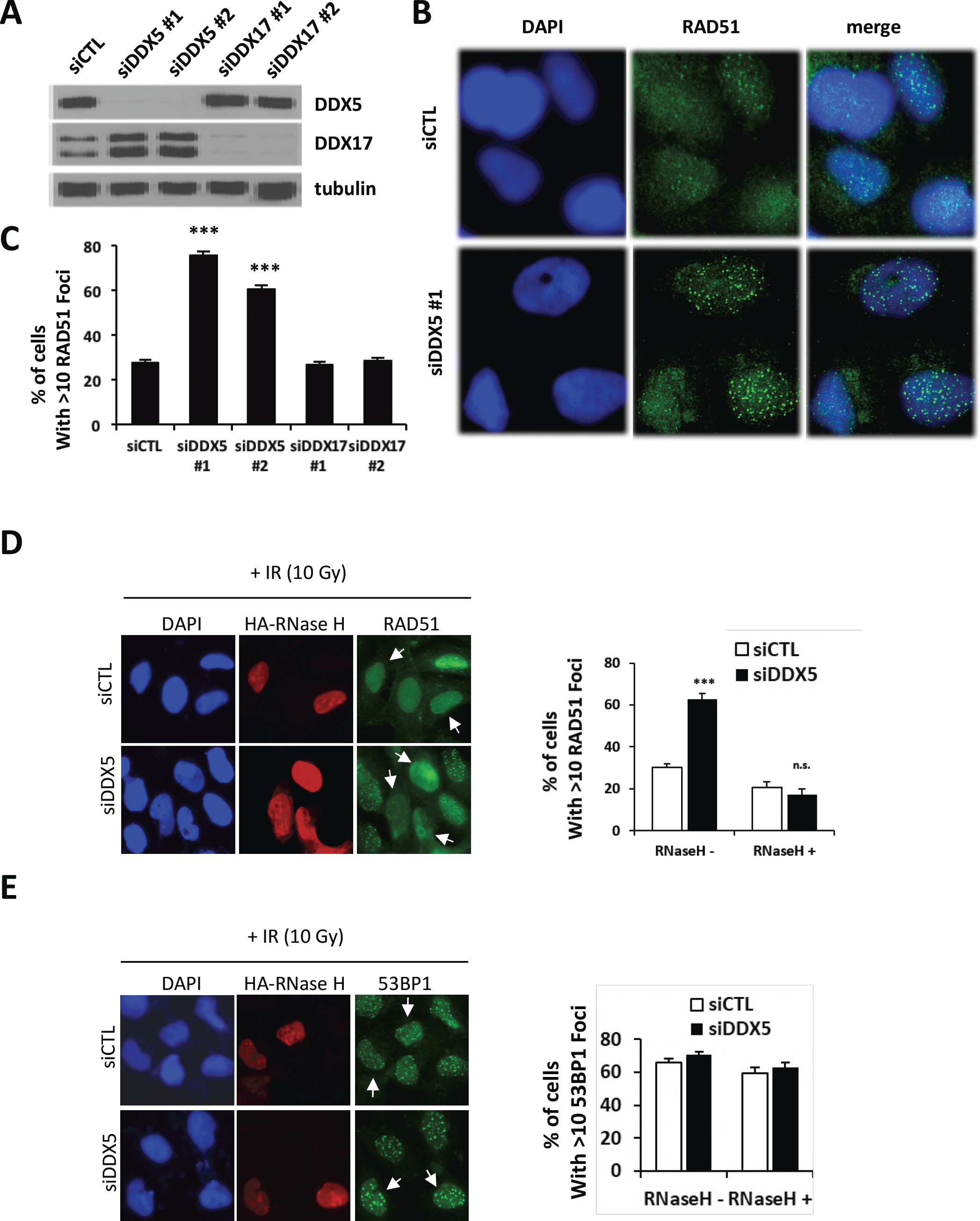
DDX5 knockdown leads to RNase H sensitive RAD51 foci. (A) U2OS cells were transfected with siDDX5 (#1 and #2), siDDX17 (#1 and #2) or control siRNA targeting luciferase (siCTL). A representative western blot analysis shows the knockdown of DDX5 and DDX17 doublet, p72 and p82 generated from alternative translation sites. (B) siCTL and siDDX5 cells were treated with 10 Gy of IR followed by 4 h recovery and visualized by indirect immunofluorescence with anti-RAD51 antibody. (C) The cells with >10 foci were counted and expressed as a percentage. The graph shows the average and standard error of the mean (S.E.M.) from two independent experiments performed in duplicates where >20 images were acquired containing at least 400 cells per condition. (D, E) U2OS cells were first transfected with control or siDDX5 siRNA and then the HA-tagged RNase H1 one day after the siRNA transfection. Two days after plasmid transfection, the cells were treated with 10 Gy of IR. After 4h of recovery, the cells were visualized by indirect immunofluorescence with anti-HA (red) and anti-RAD51 or anti-53BP1 (green) antibodies. The cells with >10 foci in each group were counted and expressed as a percentage in the group. The graph shows the average and standard error of the mean (S.E.M.) from three independent experiments where >30 different fields were analyzed. Statistical significance was assessed using Student’s t-test. *, *p* < 0.05; ***, *p* < 0.001 and *n.s*., no significant.

Many RNA helicases have been identified to regulate R-loops and it is known that a proportion of RAD51 foci are sensitive to RNaseH treatment (Wahba, Gore et al., 2013), as they are preferentially localized in transcriptionally active regions (Aymard et al., 2017). To test whether the defect of DDR in DDX5-deficient cells is associated with DDX5 function in the resolution of RNA:DNA hybrids, we investigated the effect of RNase H1 overexpression on IR-induced RAD51 foci formation. RNase H1 digests the RNA strand in the RNA:DNA hybrid, revealing R-loop dependent RAD51 foci (Wahba et al., 2013). Following siRNA control or DDX5 knockdown, U2OS cells were transfected with HA-tagged RNase H1 and visualized by indirect immunofluorescence with anti-HA (red) and anti-RAD51 or anti-53BP1 (green) antibodies. The data was analyzed in both HA-RNase H1-positive and negative populations, respectively. In the HA-RNase H1-negative population (RNase H1-), DDX5 depletion led to significant increase of the percentage of cells with more than 10 RAD51 foci, but not 53BP1 foci (Figure 1D and 1E). However, in the HA-RNase H1-transfected cells (RNase H1+), DDX5 depletion had no effect on RAD51 foci compared to control siRNA-transfected cells (Figure 1D and 1E, siCTL, RNase H1+). These results suggest that the defect of DSB response and repair in DDX5-deficient cells is associated with increased RNA:DNA hybrid formation.

### DDX5 resolves R-loops *in vitro* and *in vivo*.

DDX5 is an RGG/RG motif containing helicase previously shown to unwind RNA/RNA and RNA:DNA duplexes (Hirling, Scheffner et al., 1989, Rossler, Straka et al., 2001, Xing, Wang et al., 2017). Dbp2, the *S. cerevisiae* homolog of DDX5, was shown to resolve RNA:DNA hybrids in the context of R-loops (Cloutier, Wang et al., 2016), however, whether DDX5 shares this R-loop resolving activity is unknown. To examine whether DDX5 resolves R-loops, we purified recombinant DDX5 to homogeneity (Figure 2A) and performed *in vitro* R-loop (RNA:DNA hybrid) and D-loop (DNA:DNA hybrid) unwinding activity assays using radiolabeled nucleotide substrates. The addition of increasing DDX5 doses led to the appearance of a faster migrating species on native gels representing the DNA strands without the bound RNA fragment and this occurred in an ATP-dependent manner (Figure 2B). Remarkably, DDX5 did not resolve the D-loop substrate (Figure 2B). These observations show that DDX5, like its yeast homolog Dbp2, resolves R-loops *in vitro*.

**Figure.2.**
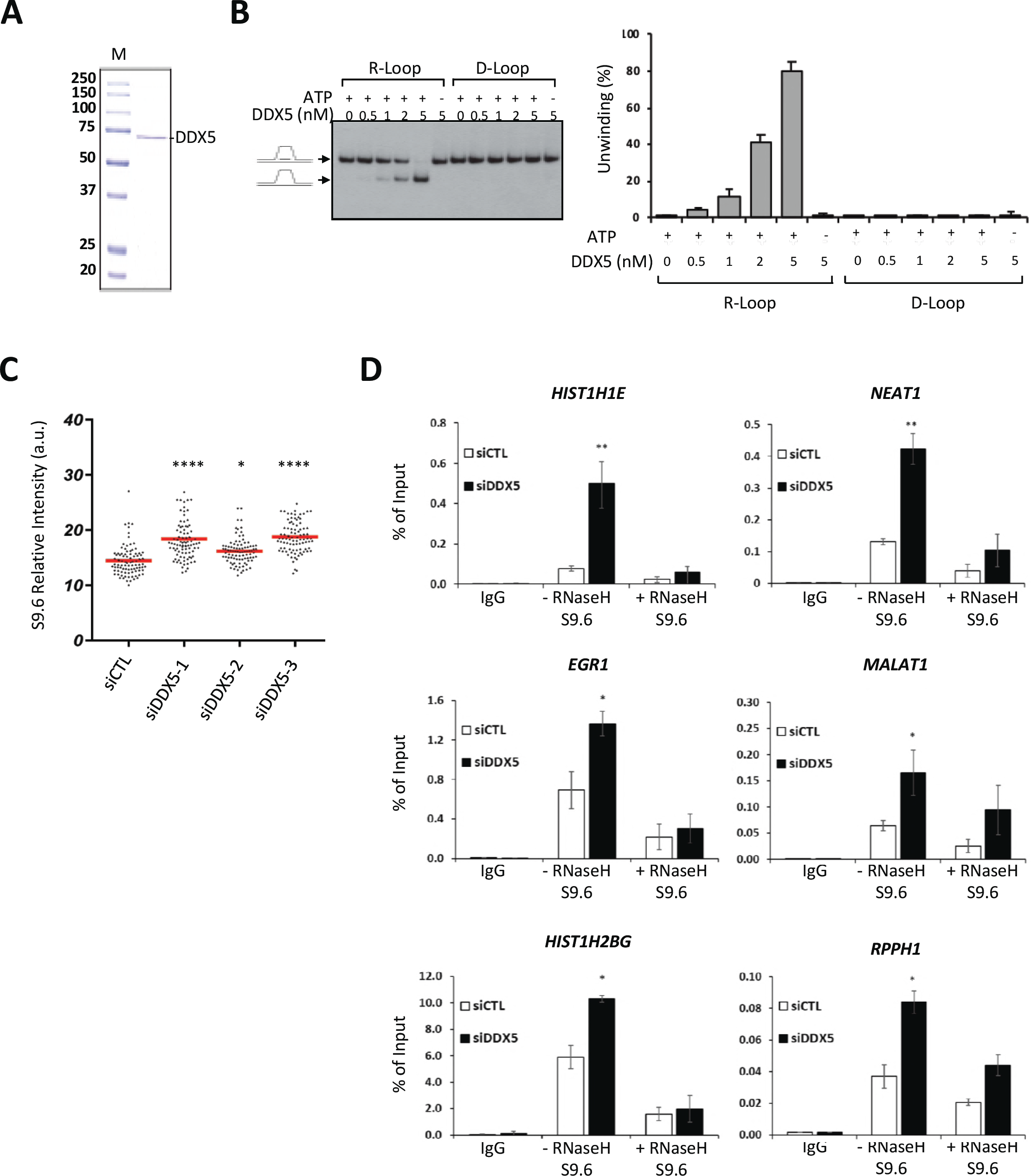
DDX5 unwinds R-loops *in vitro* and repress cellular R-loop accumulation. (A) Coomassie Blue staining of recombinant human DDX5 purified in bacteria. M denotes the molecular mass markers in kDa. (B) R-loop unwinding assay in presence of increasing DDX5. The left panel shows a typical image obtained after the assay. The bar graph (right panel) shows the quantification. The average is expressed as percentage unwinding, and standard error of the mean (S.E.M.), n=4. (C) S9.6 signal in the nucleoplasm. An average from 3 independent experiments performed in triplicate. In total, ~90 images for each condition (10 images/slide and 9 slides for each condition) were taken and two cells per image were quantified. Statistical significance was assessed using one-way ANOVA t-test. *: *p* < 0.05; ****: *p* < 0.0001. (D) U2OS cells were transfected with control (siCTL) or siDDX5 and genomic DNA was subjected to DRIP-qPCR analysis. The graphs below show the average and S.E.M. n=3. Statistical significance was assessed using t-test. *: *p* < 0.05; **: *p* < 0.01.

To examine whether DDX5 resolves R-loops *in vivo*, we generated DDX5-deficient U2OS cells using three siRNAs and we measured the accumulation of RNA:DNA hybrids by immunofluorescence using the monoclonal antibody (S9.6) known to recognize RNA:DNA hybrids within the mitochondria, nucleoli, and the nucleoplasm (Bhatia et al., 2014, Ginno, Lott et al., 2012, Hamperl, Bocek et al., 2017, Hodroj et al., 2017, Sollier et al., 2014). We measured the S9.6 signal in the nucleoplasm as the total nuclear signal subtracting the nucleolar contribution. The nucleus and nucleolus were detected with DAPI and anti-nucleolin antibodies, respectively (Figure S3A). Depletion of DDX5 led to a significant increase of the S9.6 signal in the nucleoplasm of U2OS cells with all siRNAs tested (Figure 2C). We also visualized the accumulation of RNA:DNA hybrids from isolated genomic DNA using slot-blot analysis with the S9.6 antibody in the presence or absence of RNase H. Again, we observed a significant increase in the S9.6 signal within the genomic DNA isolated from DDX5-deficient cells compared to control siRNA treated cells. siSenataxin (siSTEX) was used as a positive control (Figure S3B, S3C).

**Figure S3.**
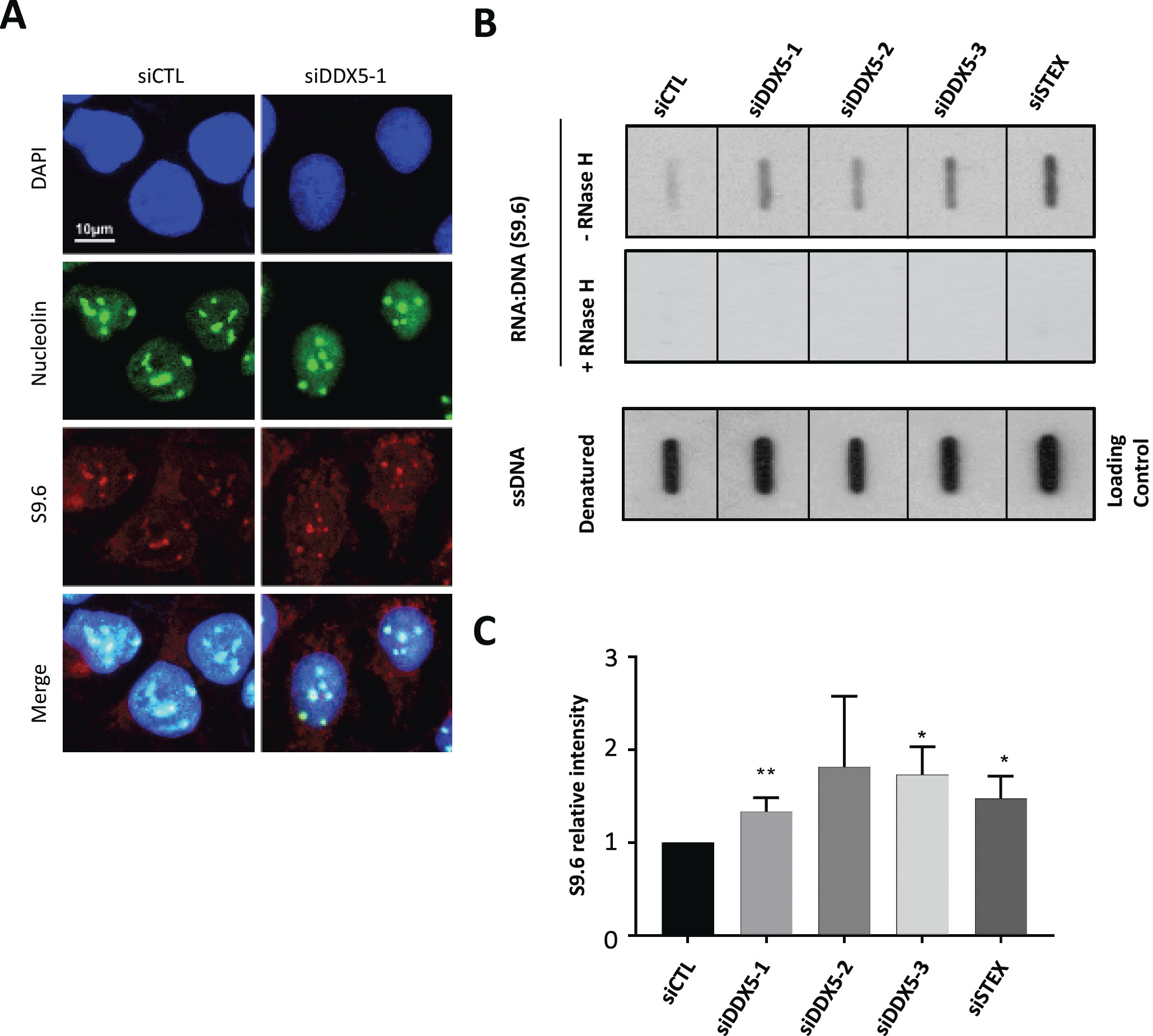
DDX5 siRNAs increase global RNA:DNA hybrids. (A) The cells were subjected to immunofluorescence analysis with S9.6 and anti-nucleolin antibodies. A typical image was shown. The S9.6 signal in the nucleoplasm is measured as the total nuclear signal subtracting the nucleolar contribution detected as nucleolin-positive areas. (B) Slot blot analysis of U2OS genomic DNA transfected with indicated siRNAs. The same amount of DNA extracted from siCTL, siDDX5, and siSTEX were left untreated, treated with RNase H (+) or denatured, as indicated. The membrane was subjected to immunoblotting with S9.6 antibodies or antibody recognizing single strand DNA (ssDNA) to verify equivalent DNA loading. (C) The graphs show the relative quantification of panel B. Statistical significance was assessed using Student’s t-test. *, *p* < 0.05; **, *p* < 0.01.

To examine the role of the DDX5 in R-loop resolution at precise genomic loci, we used the RNA:DNA immunoprecipitation (DRIP) method. By qPCR analysis we quantified R-loop accumulation at selected six specific loci: *HIST1H1E, NEAT1, RPPH1, EGR1, MALAT1*and *HIST1H2BG*, previously known to have a propensity to form R-loops (García-Rubio et al., 2015, Hodroj et al., 2017, Yang et al., 2014). The knockdown of DDX5 resulted in a significant increase of R-loops when compared to control siRNA conditions and these were sensitive to RNase H treatment (Figure 2D). These findings suggest that DDX5 deficient cells accumulate R-loops.

### DDX5 deficiency leads to spontaneous DNA damage and hypersensitivity of U2OS cells to replication stress.

As unresolved R-loops result in DNA damage (Skourti-Stathaki et al., 2011), we next investigated whether DDX5-deficient cells exhibit spontaneous DNA damage due to unresolved R-loops and indeed this was the case, as assessed by an increase in phosphorylation and foci formation of histone variant H2AX (termed gH2AX), a marker of DNA damage (Figure S4A and S4B). Cells deficient in DDX5 are known to be sensitive to ionizing radiation (Nicol, Bray et al., 2013), however it is unknown whether these cells are sensitive to DNA replication stress such as hydroxyurea, as would be predictive of R-loop accumulation during DNA replication. Indeed, depletion of DDX5 using two different siRNAs led to a significant reduction of cell survival to that of hydroxyurea treatment compared with the control siRNA-transfected cells (Figure S4C). To further confirm this effect, we also performed a FACS (fluorescence-activated cell sorting)-based survival analysis of co-cultured cells (Figure S4D). This analysis enables direct comparison in the same culture of the proliferative fitness of DDX5-expressing and depleted cells. The U2OS cells without or with stable GFP expression were transfected with control and DDX5 siRNAs, respectively. The cells were mixed and co-plated two days after transfection and then treated with hydroxyurea. Compared to non-treated cells, a decrease in the percentage of GFP-positive cells after treatment indicates an increase in sensitivity to hydroxyurea in the target siDDX5-transfected cells (Figure S4E). Depletion of DDX5 caused significant increase in cell sensitivity to hydroxyurea, either at low dose (0.4 mM) with long-time incubation (24h) or at high dose (10 mM) and short exposure (2h; Figure S4F, S4G). Taken together, these results suggest that DDX5 deficiency causes cell hypersensitivity to the DNA replication stress agent hydroxyurea.

**Figure S4.**
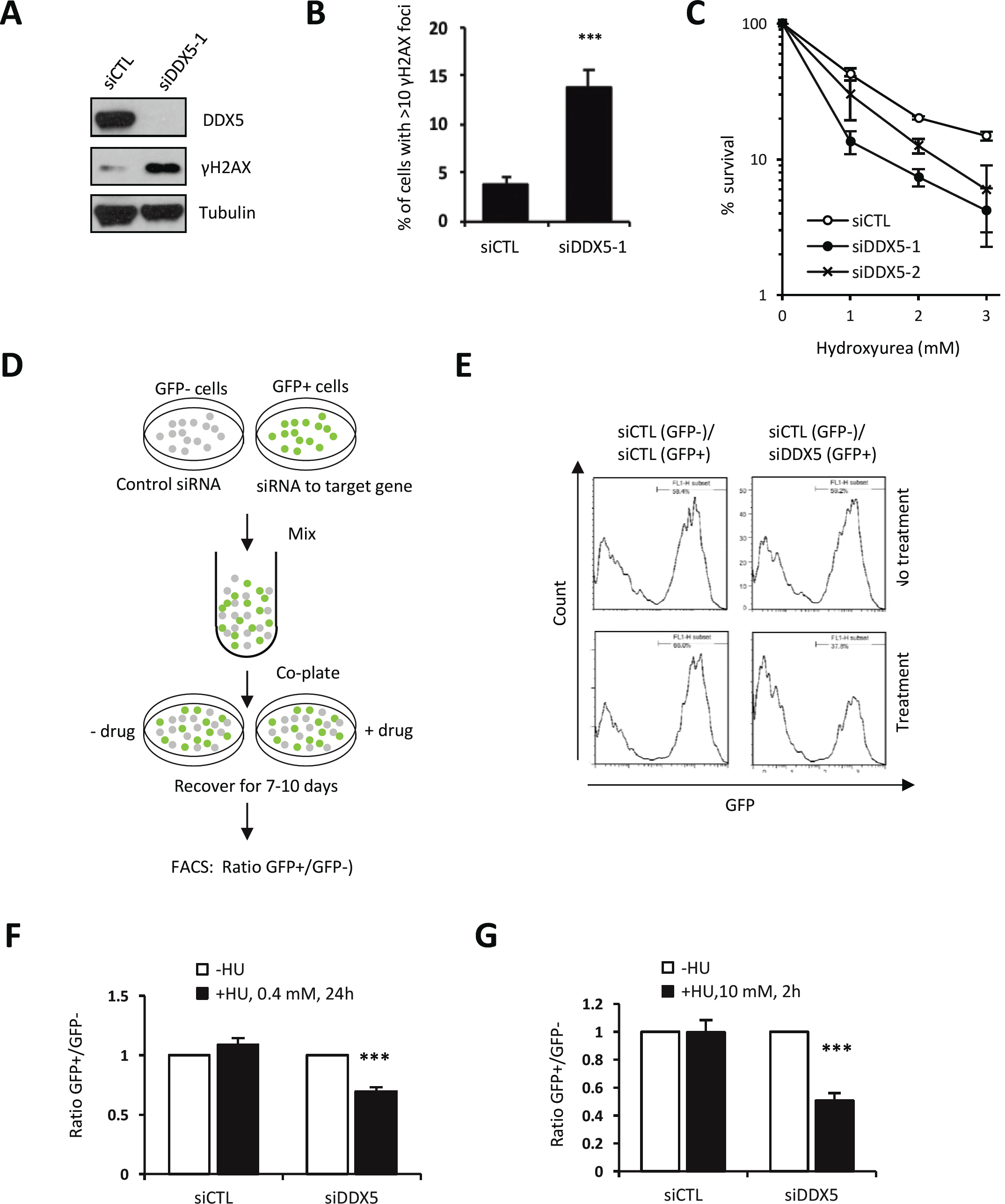
DDX5 deficiency leads to spontaneous DNA damage and hypersensitivity of U2OS cells to replication stress. (A) Immunoblotting analysis of total protein extracts derived from U2OS cells transfected with siCTL or siDDX5-1. (B) Foci quantification results after immunofluorescence with anti-γH2AX antibody of U2OS cells transfected with control (siCTL) or siDDX5-1. The cells with >10 foci were counted and expressed as a percentage of total cells. The graph shows the average and standard error of the mean (S.E.M.) from three independent experiments where >30 images were acquired containing at least 300 cells per condition. (C) Colony survival analysis of U2OS cells transfected with siDDX5 or control siRNAs. Cells were treated with different dosage of hydroxyurea (HU) for 20 h. Colony survival analysis was performed as described in the “Materials and Methods” section. The graph shows the average and standard error of the mean (SEM) from four independent experiments. (D) Illustration of FACS-based cell survival analysis. Wild type U2OS and U2OS stably expressing transfected GFP were transfected with siCTL and siDDX5. The cells were then harvested two days after transfection and mixed in an approximately 1:1 ratio. These cells were plated and exposed to hydroxyurea. Then FACS analysis was performed following seven to ten days of recovery. (E) A FACS profile shows DDX5 deficiency leads to a decrease of GFP+ cell population and an increase of GFP-cell population in the hydroxyurea-treated cells, compared to untreated cells. (F and G) Cell survival following treatment with 10 mM hydroxyurea (HU) for 2 h or 0.4 mM 24 h. Data are plotted as a ratio in GFP+ to GFP-cells normalized to untreated control. The graphs show the average and standard error of the mean (SEM) from four independent experiments. Statistical significance was assessed using Student’s t-test; ***, *p* < 0.001.

### DDX5 is arginine-methylated in U2OS cells

Interestingly, in a separate study, we identified DDX5 by mass spectrometry analysis as a PRMT5-interacting protein in two prostate cancer cell lines (Table S1). PRMT5 is known to catalyze the mono-and symmetrical dimethylation of arginine residues in proteins (Branscombe, Frankel et al., 2001). We first confirmed the physical association between PRMT5 and DDX5 in U2OS cells. Cells were lysed and either PRMT5 or DDX5 was immunoprecipitated followed by immunoblotting. Significant amounts of DDX5 was co-immunoprecipitated with anti-PRMT5 antibodies, but not control IgG (Figure 3A). The reverse was also observed, PRMT5 was coimmunoprecipitated with anti-DDX5 antibodies (Figure 3B). These findings confirm the association of PRMT5 with DDX5.

**Figure.3.**
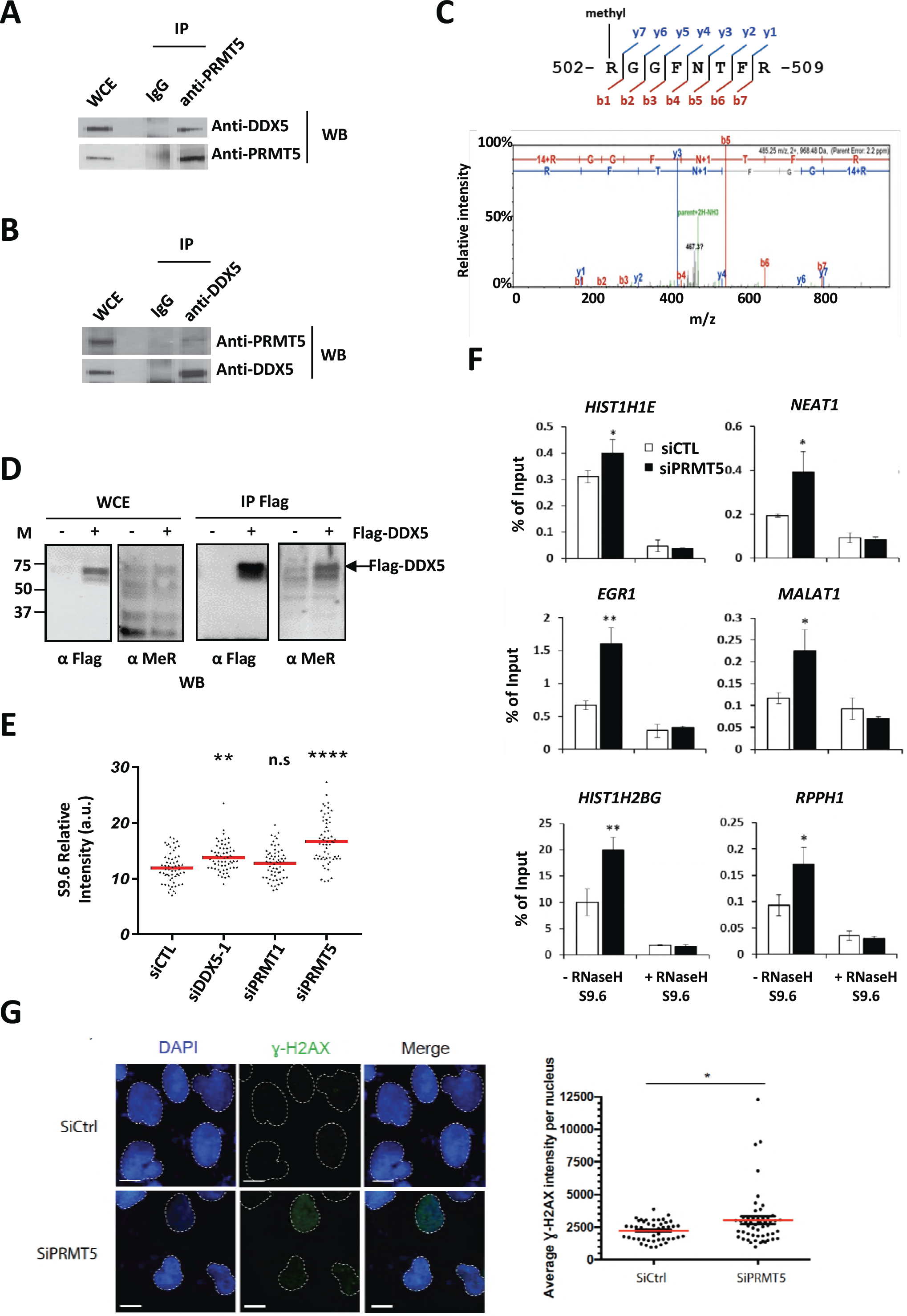
DDX5 is arginine-methylated in U2OS cells and PRMT5 deficiency causes cellular R-loops to accumulate. (A) U2OS cells were lysed and incubated with anti-PRMT5 antibody or immunoglobulin G (IgG) as control. The immunoprecipitated proteins or whole cell extract (WCE) were separated by SDS-PAGE and Western blotted (WB) with the indicated antibodies. (B) Same as panel A except anti-DDX5 antibody was used instead of anti-PRMT5 antibody for immunoprecipitation. (C) Mass spectrum shows R502 monomethylation from Flag-DDX5 expressed in U2OS. (D) HEK293 transfected with Flag-tagged DDX5 were lysed and immunoprecipitated with anti-Flag antibodies. The bound proteins were immunoblotted as indicated along with WCE from untransfected (-) and Flag-DDX5 (+) transfected cells. (E) U2OS cells transfected with the indicated siRNAs were subjected to immunofluorescence analysis with S9.6 and anti-nucleolin antibodies. Graphs show the average of 3 independent experiments performed in triplicates. (F) Genomic DNA was isolated from U2OS transfected with control (siCTL) and siPRMT5 and subjected to DRIP-qPCR. The average and S.E.M. from 3 independent experiments is shown. Statistical significance was assessed using Student’s t-test. *: *p* < 0.05; **: *p* < 0.01 ****: *p* < 0.0001. *n.s*: not significant. (G) U2OS cells were transfected with control or siPRMT5 siRNA. 72 hours later the cells were stained with anti-gH2AX (green) antibody and DAPI then visualized by indirect immunofluorescence using Zeiss LSM800 confocal system. Intensity of nuclear staining was measured in at least 50 cells chosen randomly from at least 13 different fields per condition. Each dot in the scatter plot represents the average intensity per nucleus, and the bar represents the overall average intensity per condition. Statistical significance was assessed using Student’s t-test. *, *p* < 0.05.

Next, to determine whether DDX5 is arginine methylated, we first immunoprecipitated DDX5 from U2OS cells and analyzed it for the presence of methylarginines by mass spectrometry. We identified R502 within the C-terminal RGG/RG motif of DDX5 to harbor a monomethyl (Figure 3C). U2OS cells transfected with Flag-epitope tagged DDX5 were immunoprecipitated with anti-Flag antibodies and the bound proteins immunoblotted with anti-methylarginine antibodies. We detected that Flag-DDX5 was arginine methylated (Figure 3D).

RGG/RG motifs are methylated mainly by PRMT1 and PRMT5, the two major enzymes responsible for cellular protein arginine methylation (Blanc & Richard, 2017). To identify which PRMT leads to R-loop accumulations, we turned to R-loop visualization using immunofluorescence assays with S9.6 antibody in siPRMT1 or siPRMT5 U2OS cells. We observed that the knockdown of PRMT5, but not PRMT1, led to significant increase of nuclear S9.6 staining in U2OS cells (Figure 3E). DDX5 siRNA-1 was used as a positive control. Similarly, DRIP-qPCR analysis demonstrated that depletion of PRMT5 caused significant RNA:DNA hybrid accumulation, as observed with siDDX5 (Figure 3F) and this lead to a significant increase in gH2AX foci (Figure 3G).

We next performed *in vitro* arginine methylation analysis using glutathione-S-transferase (GST)-DDX5 fusion proteins to identify the methylation region. DDX5 has two RGG/RG motifs: One located at its N-terminus and the other at its C-terminus. Interestingly, both DDX5 RGG/RG motifs are conserved in the yeast homolog Dbp2, although the rest of the C-terminal region is truncated in Dbp2 (Figure 4A). We generated three truncation mutants of DDX5, including the N-terminal region (residues 1-100; F1), the central catalytic enzyme domain (92-471; F2) and the C-terminal region (residues 466-614; F3; Figure 4B). Only the C-terminal region (F3), encompassing the RGG/RG motif, was methylated by PRMT5 (Figure 4C). We then substituted DDX5 R478, R482, R484, R486, R502 within the RGG/RG motif with lysines in a smaller region 466-555; F4). The 5R to 5K mutation within DDX5 (RK) completely abolished DDX5 arginine methylation by PRMT5 (Figure 4D).

**Figure.4.**
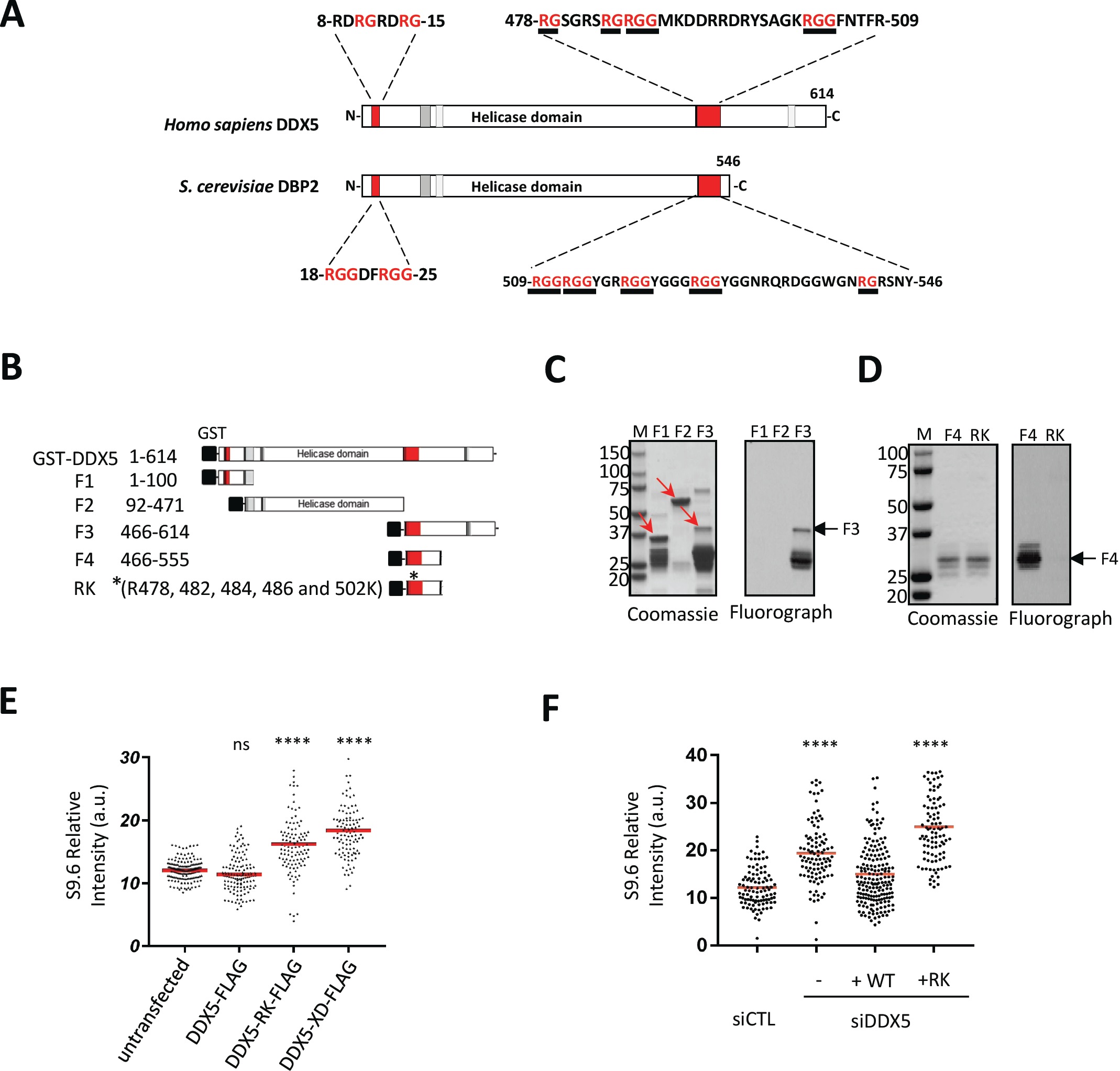
The RGG/RG motif of DDX5 is a substrate for PRMT5 and required for DDX5 function in R-loop resolution. (A) A schematic of DDX5 functional domains and motifs is shown. (B) GST-fusion of DDX5 fragments (F1-F4) and the RK mutant used in this study. RK (*) indicates R-K substitutions introduced of 5 arginines (R:478,482,484,486,502:K). (C and D) Coomassie blue (left panel) and *in vitro* methylation assay (right panel) of indicated GST-DDX5 fragments and the RK mutant. (E) Immunofluorescence analysis with S9.6 and anti-Flag antibodies of U2OS cells transfected with Flag-tagged DDX5, DDX5-RK or DDX5-XD (helicase dead). Nuclear S9.6 signal was counted in both Flag-negative and Flag-positive cells. The Flag-negative cells were considered as untransfected cells. The graphs shown represent the quantification with the S.E.M. (F) Immunofluorescence analysis with S9.6 and anti-Flag antibodies of U2OS cells transfected with siCTL or siDDX5-1 and Flag-tagged DDX5 or DDX5-RK (+RK) as indicated. The graphs show the average S.E.M. Statistical significance was assessed using one-way ANOVA t-test. ****, *p* < 0.0001.

### DDX5 helicase activity and RGG/RG motif is required for R-loop resolution.

Next, we examined whether R to K substitutions of DDX5 affected its R-loop resolution function. Interestingly, ectopic expression of DDX5-RK as well as the catalytically inactive mutant (DDX5-XD) caused significant increase of R-loop accumulation in U2OS cells, as determined by S9.6 immunofluorescence (Figure 4E). These findings suggest that the ectopic expression of DDX5-RK and DDX5-XD likely behave as dominant negatives. We further investigated whether WT DDX5, DDX5-RK, and DDX5-XD can rescue the R-loop accumulation observed in DDX5 deficiency. The ectopic expression of WT DDX5 reversed the R-loop accumulation observed in siDDX5 cells, however, this was not observed with the expression of DDX5-RK (Figure 4F). Taken together, these results suggest that the RGG/RG motif is required for R-loop resolution.

### DDX5 associates with known R-loop regulatory proteins.

To identify interacting partners of DDX5 which may function in R-loop metabolism, we performed stable isotope labeling with amino acids in cell culture (SILAC)-based mass spectrometry analysis. U2OS cells expressing Flag-epitope tagged DDX5 were grown in the ‘heavy’ mediurm and the control U2OS cells in light medium (Figure S5A). Many RNA binding proteins belonging to the heterogeneous nuclear ribonucleoproteins (hnRNPs) and DEAD/DEAH-box families were identified (Table S2 and Figure S5B). These included known DDX5 interactors including DHX9, DDX3X, and DDX17 (Choi & Lee, 2012, Ogilvie, Wilson et al., 2003, Wilson & Giguere, 2007). Interestingly, we also identified proteins known to influence R-loops resolution: DDX1, DHX9, XRN2, SRSF1 and Aly/REF an exon junction complex protein (Table S2). PRMT5 was also identified, but since PRMT5 is a contaminant of anti-FLAG immunoprecipitates, we could not claim association with DDX5 based on this approach.

**Figure S5.**
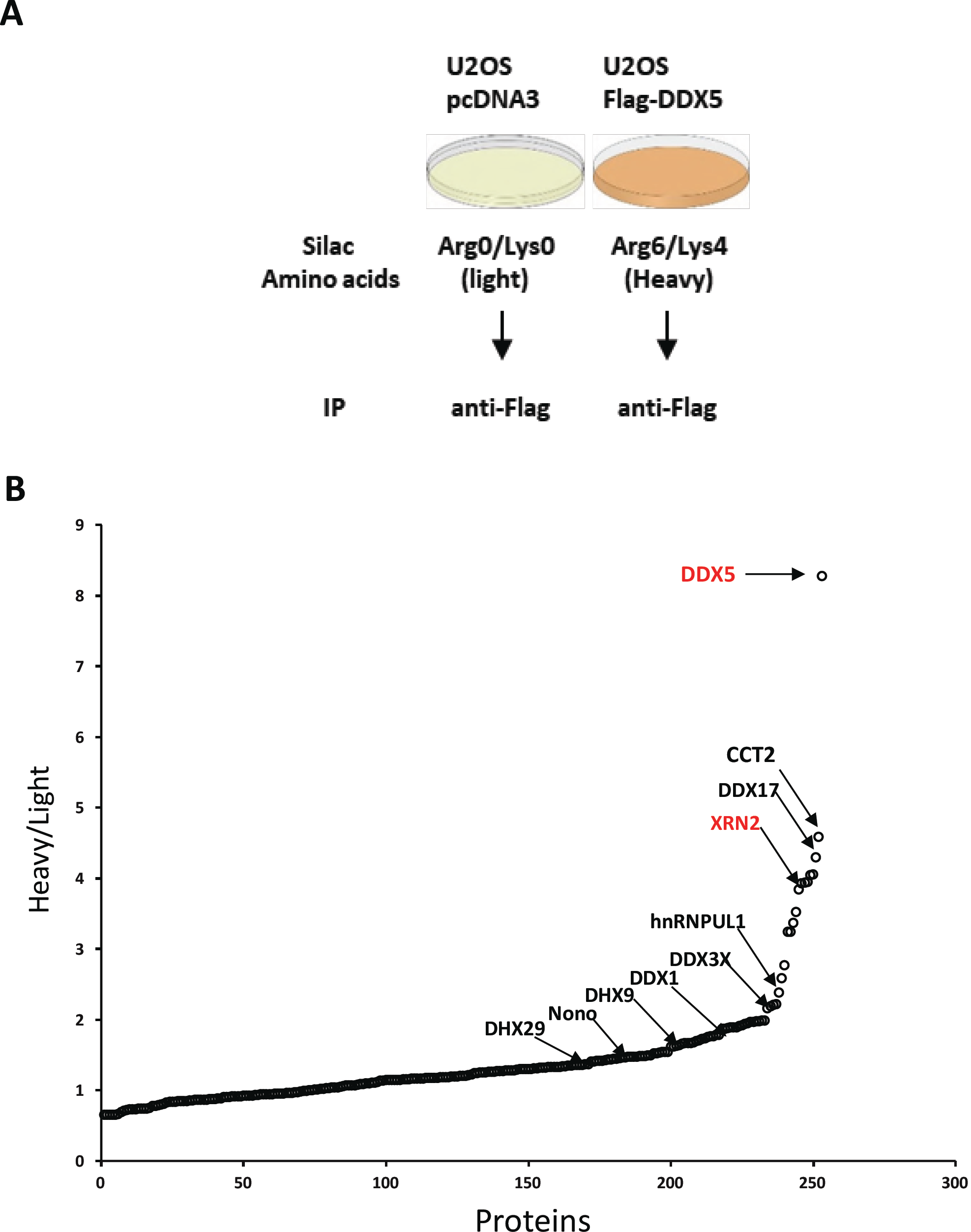
Identification of DDX5-interacting proteins by SILAC MS/MS spectrometry. (A) U2OS cells with stably or transiently transfected Flag-DDX5 or empty vector were subjected to SILAC analysis as described in the “Materials and Methods” section. (B) The identified proteins, including XRN2 with a heavy/light signal ratio were presented. (XRN2 ratio is 3.92).

### RGG/RG motif of DDX5 mediates association with XRN2

XRN2 has been shown to function with helicases such as Senataxin (Cristini et al., 2018, Skourti-Stathaki et al., 2011) and DHX9 (Cristini et al., 2018) and is required to resolve R-loops (Morales et al., 2016). We hypothesized that XRN2 also assists DDX5 with R-loop resolution of a subset of genes. We first confirmed the interaction between the two proteins. U2OS cells expressing Flag-DDX5, were immunoprecipitated with anti-Flag antibodies and the presence of XRN2 was detected by immunoblotting (Figure 5A), confirming our mass spectrometry data. Endogenous DDX5 also co-immunoprecipitated with XRN2, but not control IgG (Figure 5B). We next mapped the region of DDX5 required to associate with XRN2. DDX5 is composed of a central helicase domain with N-and C-terminal RGG/RG motifs (Figure 4A). The XRN2 interaction region was mapped between amino acids 435 and 554 of DDX5, where the C-terminal RGG/RG motif resides (Figure 5C). We next determined whether mutation of these arginine residues within the RGG/RG motif affected recognition with anti-methylarginine antibody and interaction with XRN2. Indeed, Flag-DDX5:RK was not recognized with the anti-methylarginine antibody and had reduced association with XRN2, unlike wild type DDX5 (Figure 5D). These findings suggest that arginine methylation of the RGG/RG motif of DDX5 regulates the interaction with XRN2.

**Figure.5.**
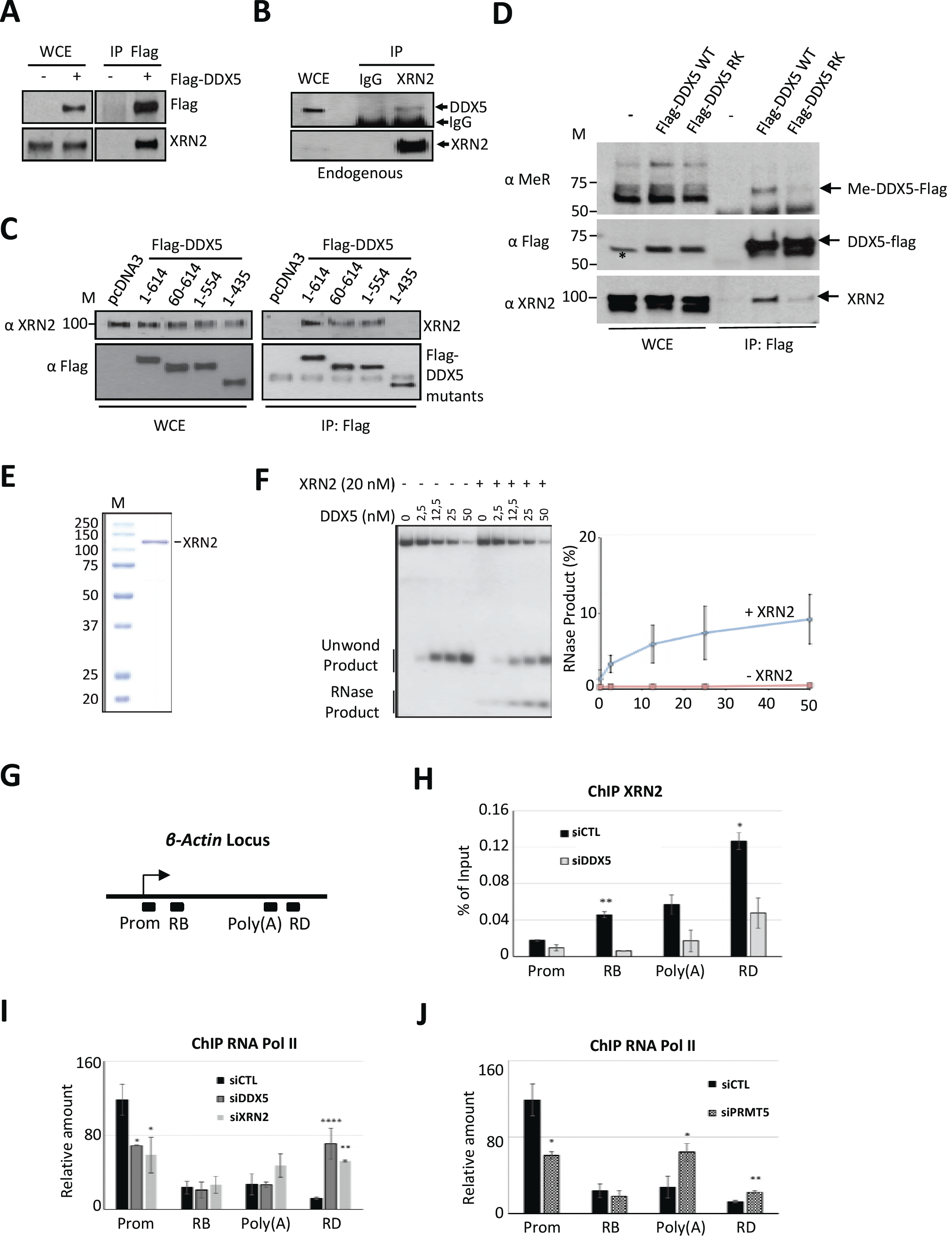
DDX5 physically associates with the 5’-exonuclease XRN2, functioning together to repress R-loops. (A) HEK293 cells were transfected with Flag-DDX5 (+) or empty pcDNA3 vector (-). Whole cell lysates (WCE) were subjected to immunoprecipitation with anti-Flag antibody and immunoblotting with indicated antibodies. (B) U2OS lysates were subjected to immunoprecipitation with control immunoglobulin G (IgG) or anti-XRN2 antibodies and immunoprecipitated proteins separated by SDS-PAGE followed by immunoblotting with the indicated antibodies. (C) U2OS cells were transfected with expression vectors that encode Flag-DDX5 full-length or truncated mutant proteins. Anti-Flag immunoprecipitations were monitored for the presence of endogenous XRN2 by immunoblotting with anti-XRN2 antibodies. (D) U2OS cells transfected with Flag-tagged DDX5 were subjected to immunoprecipitation with anti-Flag antibody and immunoblotted with anti-Flag, anti-XRN2 and methylarginine-specific (Me-R) antibody, respectively. (E) Coomassie Blue staining of recombinant human XRN2 purified from bacteria. M indicates the molecular mass markers in kDa. (F) R-loop unwinding and RNA exonuclease activity analysis. A typical image is shown on the left. The data was analyzed as percentage of RNase product (right panel). The graph show the average and S.E.M. n=4. (G) Schematic representation of the b*-actin* locus and the location of the PCR primers used for ChIP-qPCR analysis. Prom (promotor), RB (region B), poly(A) and RD (region D) indicated the region analyzed by qPCR. (H) HEK293 cells were transfected with siCTL or siDDX5. The XRN2 ChIP signal normalized to the Pol II ChIP signal with or without DDX5 depletion is shown. (I) ChIP analysis of RNA Pol II on the b-Actin locus in cells transfected wit siCTL, siDDX5 or siXRN2. The Y-axis shows the signal-to-noise ratio of RNA Pol II IP relative to control IP. (J) ChIP analysis of RNA Pol II on the b-actin locus after siCTL or siPRMT5. The Y-axis shows the signal-to-noise ratio of RNA Pol II IP relative to control IP. The graphs show the average and S.E.M. Statistical significance was assessed using Student’s t-test. * *p* < 0.05, ** *p* < 0.01 and **** *p* < 0.0001.

### DDX5 and XRN2 function together in R-loop resolution

To test whether DDX5 and XRN2 function together in R-loop resolution, we performed DDX5 R-loop resolving *in vitro* assays in the presence or absence of purified XRN2 (Figure 5E). The presence of recombinant XRN2 (20 nM) increased the RNase product (i.e. degraded RNA) component of the RNA:DNA hybrid unwound by DDX5 (Figure 5F). However, XRN2 did not influence the overall ability of DDX5 to resolve R-loops, as observed by the generation of the unwound product which was the same in the absence or presence of XRN2 (Figure 5F). These results show that XRN2 degrades the RNA released by DDX5 *in vitro*.

The activity of the XRN2 exonuclease is required to promote transcriptional termination of a subset of genes (Kim, Krogan et al., 2004, West, Gromak et al., 2004). Given the requirement of XRN2 for efficient termination, we examined whether DDX5 might play a role in coupling R-loops resolution for transcription termination of certain genes. To test this possibility, we performed ChIP assays to measure XRN2 occupation along the *β-actin* gene in the DDX5 depleted cells. HEK293 cells transfected with siCTL or siDDX5 showed XRN2 distribution throughout *β-actin*, as previously shown (Kaneko, Rozenblatt-Rosen et al., 2007). However, a significant decrease of XRN2 occupancy was observed in siDDX5 depleted cells (Figure 5G, H; region B, poly(A) and region D). These findings suggest that DDX5 can regulate the recruitment of XRN2 at *β-actin* locus. To test whether this effect is potentially associated with transcriptional termination, we performed ChIP to detect the occupancy of the RNA polymerase II (PolII) downstream of the poly(A) site at the termination region. DDX5 depleted cells showed a dramatic increase in RNA PolII deposition at the *β-actin* terminator region, as observed in XRN2 depleted cells (Figure 5I, region D). These results suggest that DDX5, like XRN2 plays a role in the vicinity of the poly(A) site to ensure the release of RNA PolII for termination. We next examined whether siPRMT5 depleted cells also harbored an increase in RNA PolII recruitment at the termination region. Indeed, ChIP analysis showed an increase occupancy of RNA PolII in the absence of PRMT5 at the poly(A) and region D (Figure 5J). These observations show that PRMT5 deficiency increases RNA PolII increase at the *β-actin* transcription termination site consistent with R-loop accumulation, as shown previously (Zhao et al., 2016).

## Discussion

In this study, we define a new function for DDX5 in R-loop resolution and reveal a regulatory role for the methylation of its RGG/RG motif in modulating its function. Depletion of either DDX5 or PRMT5 *in cellulo* lead to increased accumulation of RNA:DNA hybrids, as detected with the anti-S9.6 antibody by immunofluorescence, slot blot analysis and DRIP-qPCR. The C-terminal RGG/RG motif of DDX5 was methylated by PRMT5 and required for R-loop resolution, as amino acid substitution of its arginines in the RGG/RG motif with lysines (RK) failed to reverse the accumulation of R-loops in siDDX5 cells. Purified human DDX5 resolved R-loop structure *in vitro* and its catalytically inactive mutant failed to rescue the effect of DDX5 deficiency on R-loop accumulation, suggesting that DDX5 represses R-loop accumulation by directly resolving the RNA:DNA hybrid structures. Moreover, proteomic analysis revealed that DDX5 associates with many RNA binding proteins including helicases, as well as with XRN2, an exonuclease known to function in transcriptional termination (Kim et al., 2004, West et al., 2004). This is in line with a few recent studies for high-through-put identification of R-loop associated proteins, in which DDX5 was identified as a top hit of R-loop associated proteins (Cristini et al., 2018, Wang, Grunseich et al., 2018), although DDX5 function was not addressed in these studies. We show that one mechanism of action of DDX5 is to mediate regulated interaction with XRN2. DDX5-deficient cells harbored less XRN2 at the b*-actin* transcriptional termination site, accompanied with an increase of RNA pol II accumulation. These data are consistent with pausing and R-loop accumulation and DDX5 functioning to recruit XRN2 at certain genomic regions in a PRMT5 dependent manner. It is known that p54nrb and PSF function to recruit XRN2 at transcription termination sites (Kaneko et al., 2007), therefore, DDX5 may function with these factors for optimal XRN2 recruitment.

Consistent with DDX5 function in R-loop repression, DDX5 knockdown lead to an increase number and a sustained presence of RNase H-sensitive RAD51 foci in response to IR-induced DSBs. Recently, a few studies suggest that HR proteins, such as RAD51, RAD52 and RPA1 are selectively recruited to transcriptionally active DNA damage sites (Aymard et al., 2017). Genome-wide mapping reveals that RNA:DNA hybrids accumulate at DSB sites and Senataxin can remove DSB-induced RNA:DNA hybrids (Cohen, Puget et al., 2018). The increase of IR-induced RAD51 foci in DDX5-deficient cells is likely associated with an increase of RNA:DNA hybrid accumulation surrounding DSBs as DDX5 has a similar function as Senataxin for R-loop removal. Transcription-associated R-loop formation at DSBs frees non-template single-stranded DNA (ssDNA) which may facilitate recruitment of ssDNA-binding proteins such as RPA and RAD51 to DSBs and HR repair. Alternatively, R-loop removal and formation at DSBs may be associated with DSB-induced transcription silencing (Shanbhag, Rafalska-Metcalf et al., 2010) or activation (Francia, Michelini et al., 2012), which in turn determine DSB repair pathway choice. For example, DSB-induced transcription silencing may facilitate NHEJ protein recruitment and DNA end ligation, and the transcription activation may facilitate RNA:DNA accumulation and HR protein recruitment as well as RNA-mediated HR repair.

The methylation of the RGG/RG motif of DDX5 likely serves additional functions besides recruiting XRN2 and this is especially pertinent in R-loops that are not associated with transcriptional termination regions. Furthermore, the function we ascribe to the methylation of DDX5 probably functions in a coordinate fashion with the methylation of other substrates such as RNA polII subunit POLR2A to regulate R-loops suppression (Zhao et al., 2016). Besides DDX5, another DEAD box helicase, DDX21 recently shown to regulate R-loops (Song et al., 2017) harbors an RGG/RG motif and may be regulated in a similar manner as DDX5. As Tudor domains are known to interact with methylated arginines (Bedford & Clarke, 2009), we do not know the protein interface that XRN2 uses to interact with methylated DDX5. Of note, XRN2 itself also contains an RGG/RG motif, however, we do not know its function.

Why so many helicases are required to resolve R-loops? Certain RNA helicases may function on different types of R-loops or regulate R-loops for a particular process or at different genomic locations. For example, DDX1 converts G4 RNA structures to R-loops for IgH class switch recombination (Ribeiro de Almeida et al., 2018). DDX1 also functions in tRNA, mRNA and microRNA processing, transport of RNAs from nucleus to cytoplasm, and AU-rich element-mediated mRNA decay in cytoplasm. Its function in R-loop resolution facilitates homologous recombination repairs (Li et al., 2016). DDX19 is well-known for its functions in mRNA nuclear export at the nuclear pore. A role for DDX19 in R-loop suppression has been reported, suggesting a functional link between mRNA nuclear export and R-loop clearance (Hodroj et al., 2017). DDX23 is a component of the spliceosomal U5 small nuclear ribonucleoprotein (U5 snRNP) and required for integration of U4/U6-U5 tri-snRNP into the splicesome. However, the role of DDX23 in suppressing R-loops does not require a functional U5 snRNP (Sridhara et al., 2017). DDX21 is known to be involved in multiple steps of ribosome biogenesis as well as mRNA transcription (Song et al., 2017).

PRMT5 deficient cells exhibit induction of the p53 response, DNA damage and cell death and as such PRMT5 is an interesting therapeutic target for many cancers. The upregulation of the p53 response was shown to be the result of an imbalance in the methylation of Sm proteins and other RNA binding proteins (e.g. SRSF) that regulate *MDM4* and *MDM2* alternative splicing (Bezzi et al., 2013). PRMT5 deficient cells also accumulate DNA damage leading to cellular senescence or cell death (Bezzi et al., 2013, Clarke et al., 2017, Hamard et al., 2018). What leads to the increased DNA damage in these cells is still not clear. Our observations herein suggest that PRMT5 defective cells fail to suppress R-loops leading to increase DNA damage and it may represent one the earliest response to PRMT5 deficiency. The DNA damage observed from the R-loop accumulation may also contribute to increasing the p53 response, as DNA damage is known to upregulate p53.

## Acknowledgements

We thank Drs. Mark Bedford and Yanzhong Yang for purified S9.6 antibody and helpful discussions. This work was funded by FDN-154303 to S.R. and FDN-388879 to J.Y.M. M.K. is a recipient of a Doctoral Cole Foundation Fellowship. F.F.B. received a scholarship from the Emerging Leaders in the Americas Program (ELAP). J.Y.M. is a FRQS chair in genome stability.

## MATERIALS AND METHODS

### Reagents and Antibodies

Mouse anti-DDX5 monoclonal antibodies (A-5, sc-166167 and clone204, 05-850) were purchased from Santa Cruz Biotechnology and Millipore. Rabbit anti-RAD51 (H-92, sc-8349) antibody was purchased from Santa Cruz Biotechnology. Rabbit anti-53BP1 antibody (NB100-304) was from Novus Biologicals. Rabbit anti-nucleolin antibody (ab50279) and mouse anti-RPA2 monoclonal antibody (ab16850) was from Abcam. Anti-BrdU (B44) monoclonal antibody was from BD Biosciences. Anti-XRN2 antibodies (A301-102A and Ab72181) were purchased from Bethyl Laboratories Inc and Abcam (ab72181) for western blot, and from Proteintech (11267-1-AP) for ChIP experiment. Mouse anti-γH2AX (05-636) monoclonal antibody and anti-PRMT5 antibody was obtained from Millipore. S9.6 antibody was a kind gift from Drs. Bedford and Yang (Yang et al., 2014) used for IF experiments and western blot. For the DRIP experiments S9.6 was purified from the hybridoma purchased from the American Type Culture Collection (ATCC, Manassas, VA). Anti-nucleolin antibody (ab50279) was purchased from Abcam. Anti RNA PolII from Santa Cruz Biotechnology, Inc (CTD4H8: sc-47701). Alexa Fluor-conjugated goat anti-rabbit antibodies and anti-mouse antibodies were from Invitrogen. *Escherichia coli* RNase H was purchased from New England Biolabs. Hydroxyurea, protein ASepharose, mouse anti-Flag and α-tubulin monoclonal antibodies were from Sigma. Protease inhibitor cocktail and protein phosphatase inhibitor cocktail were from Roche. Monomethyl arginine (R*GG) (D5A12) Rabbit mAB was from Cell Signaling.

### Cell Culture, siRNAs, Plasmids and Transfection

All mammalian cells were cultured at 37°C with 5% CO_2_. U2OS human osteosarcoma cells (ATCC) and HEK293 cells (ATCC) were cultured in Dulbecco’s modified Eagle’s medium containing 10% v/v fetal bovine serum (FBS). U2OS cells were transfected with plasmid DNAs using Lipofectamine 2000 or 3000 and siRNA oligonucleotides using Lipofectamine RNAiMAX (Invitrogen) according to the manufacturer's instructions. HEK293 cells were transfected with plasmid DNAs by standard calcium phosphate precipitation.

All siRNAs were purchased from Dharmacon. siRNA sequences are as follows: siDDX5 #1, 5’-ACA TAA AGC AAG TGA GCG AdTdT-3’; siDDX5 #2, 5’-CAC AAG AGG UGG AAA CAU AdTdT-3’; siDDX5 #3, 5’-CAA GUA GCT GCU GAA UAU UUU-3’; siXRN2, Smartpool siGENOME human XRN2 siRNA (M-017622-01); siSenataxin, Smartpool siGENOME human SETX siRNA (M-021420-01), siPRMT1, 5'-CGT CAA AGC CAA CAA GTT AUU-3'. siPRMT5, 5’-UGG CAC AAC UUC CGG ACU UUU-3’. The siRNA 5’-CGU ACG CGG AAU ACU UCG AdTdT-3’, targeting the firefly luciferase (GL2) was used as control. 20 nM siRNA was used for transfection. For co-transfection of two or more siRNAs, the total siRNA amount was adjusted to be the same in each sample by adding control siRNA (siLuc, GL2).

The N-terminal Flag-tagged DDX5 plasmid was constructed by inserting a Flag-coding sequence into the pcDNA3.1 (+) vector at the *Hind* III and *Bam* HI sites to get pcDNA3.1-Flag and then the PCR-amplified human DDX5 cDNA coding region at *Bam* HI and *Xho* I sites of pcDNA3.1-Flag vector. The Flag-DDX5 codon-silent mutant resistant to all three siDDX5 siRNAs used in this research was constructed in the pcDNA3.1-Flag vector using Gibson Assembly Cloning Kit (New England BioLabs Inc.) according to the manufacturer's instructions. The gBlock DNAs were synthesized by Integrated DNA Technologies (IDT). The catalytic inactive DDX5 mutant with replacement of both Arg403 at the motif Va and Arg428 at the motif VI with Leucine and the RK mutant with replacement of the five arginine residues with lysine at the RGG/RG motif were constructed by two-steps PCR using the siRNA-resistant DDX5 construct as template. The plasmids for expressing GST fusion proteins of DDX5 fragments were constructed by inserting PCR-amplified DDX5 cDNA fragments in pGEX-6P1 vector at *Bam*HI and *Sal* I sites. HA tagged RNase H1 was a kind gift from Dr. Yangzhong Yang (Beckman Research Institute, City of Hope, CA).

### Cell Lysis and Immunoprecipitation

For co-immunoprecipitation experiments, cells were lysed with a lysis buffer containing 50 mM HEPES, pH 7.4, 150 mM NaCl, 1% Triton X-100 and a cocktail of protease inhibitors and phosphatase inhibitors. After high speed centrifugation, the supernatant was precleared and then incubated with antibodies for one hour and then protein-A or protein G agarose beads for another hour at 4°C. In some experiments, the supernatant was incubated directly with agarose beads on which antibodies were pre-attached covalently. The beads were washed five times with lysis buffer for western blot analysis or four time with lysis buffer and then twice with PBS for mass spectrometry analysis.

### SILAC (Stable Isotope Labeling with Amino Acids in Cell Culture) and MS/MS Mass Analysis

U2OS stable cells with Flag-DDX5 expression were grown in standard heavy (Lys4/Arg6) SILAC medium and the control U2OS cells in light medium (Lys0/Arg0) for ten days. Similarly, U2OS stable cells with Flag-DDX5 expression were grown in the heavy SILAC medium and the control U2OS cells in light medium for ten days. The cells were lysed and subjected to immunoprecipitation with anti-Flag antibody as described above. The beads with bound proteins were sent as described previously (Thandapani, O'Connor et al., 2013).

### Immunofluorescence

For immunostaining with S9.6 and anti-nucleolin antibody, the cells were fixed with cold methanol for 10 minutes at room temperature. Slides were then washed three times with PBS containing 0.1% Tween-20 (PBST) and blocked with blocking buffer (3% bovine serum albumin in PBST) for 1 h at room temperature or overnight at 4**°**C. Slides were then incubated with antibodies S9.6 (1/200) and anti-nucleolin (1/1000) diluted in the blocking buffer for 2 hours. After three washes with PBST, slides were incubated with corresponding fluorescent secondary antibodies for 2 h at room temperature. Slides were then washed three times with PBST before mounting with IMMU-MOUNT (Thermo Scientific) mounting medium containing 1µg/ml of 4’, 6-diamidino-2-phenylindole (DAPI). For immunostaining with anti-γ-H2AX, RAD51 and 53BP1 antibody, cells were fixed for 10 min with 4% paraformaldehyde (PFA). For RPA2 and BrdU foci detection, cells were pre-extracted for 10 min on ice with the extraction buffer 1 (10 mM PIPES, pH 7.0, 100 mM NaCl, 1 mM EGTA, 3 mM *MgCl_2_*, 300 mM sucrose and 0.5% Triton X-100) and then 10 min with extraction buffer 2 (10 mM Tris-Cl*_2_*, pH7.5, 10 mM NaCl, 1% Tween 40 and 0.5% sodium deoxycholate) before being subjected to paraformaldehyde fixation. After three washes with PBS, the cells were permeabilized for 5 min with 0.5% Triton X-100 in PBS. Coverslips were incubated overnight in PBS blocking buffer containing 10% FBS and 0.1% Triton X-100, and then incubated with primary antibody against γ-H2AX (1:2000), 53BP1 (1:200), RAD51 (1:20), RPA2 (1:500), BrdU (1:100), Flag (1:200) or HA (1:200) antibody diluted in PBS containing 5% FBS for 30 min. After three washes, the coverslips were incubated with corresponding fluorescent secondary antibodies for another 30 min in PBS buffer containing 5% FBS. After rinsing, the coverslips were mounted with IMMU-MOUNT (Thermo Scientific) mounting medium containing 1µg/ml of 4’, 6-diamidino-2-phenylindole (DAPI). Images were taken using a Zeiss M1 fluorescence microscope with 63X amplification and Zeiss LSM800 confocal system with 40X magnification where indicated.

### Slot-blotting

Nucleic acids were extracted from U2OS cells by SDS/proteinase K treatment at 37°C overnight followed by phenol-chloroform extraction and ethanol precipitation. The nucleic acids were blotted onto Hybond-N Nylon membrane in duplicate using a Slot-Blot apparatus (Schleicher & Schuell). One half of the membrane was treated with 0.5 N NaOH, 1.5 M NaCl for 10 min to denature the DNA and neutralized for another 10 min in 0.5 M Tris-HCl buffer (pH7.0) containing 1 M NaCl. After UV-crosslinking (0.12 J/m2), the non-treated membrane was subjected to western blot analysis with S9.6, and the treated membrane, served as loading control, with the single-stranded DNA antibody. The S9.6 signal was normalized by the loading control.

### DRIP (RNA:DNA Immunoprecipitation)-qPCR

DRIP assays were performed as described (Ginno et al., 2012). Briefly, nucleic acids were extracted from U2OS cells by SDS/proteinase K treatment at 37°C overnight followed by phenol-chloroform extraction and ethanol precipitation. The harvested nucleic acids were digested for 24 h at 37°C using a restriction enzyme cocktail (50 units/100 µg nucleic acids, each corresponding to 4x10^6^ cells were resuspended in 1x Chip sonication lysis buffer in presence of protease inhibitors (PIC) and sonicated to achieve a chromatin sized of 300-1000 bp using a Branson 450 CE Sonicators (total 4 min run, 1 sec ON and 1 sec OFF) on ice. After pelleting debris, the equivalent of 10 µg of sonicated, cross-linked chromatin were incubated overnight with desired antibody and then incubated 2h with protein G magnet beads. The beads were washed with furnished buffer, as indicated by the protocol kit. Chromatin was eluted from the beads with elution buffer at 65^o^C. The cross-linking of the eluted chromatin as well as the input were treated for 30 min with RNase A at 37°C in presence of 200mM of NaCl, and then reversed cross-linked at 65 °C for 2h in presence of proteinase K. The chromatin was purified by column purification kit and eluted in 50µl of elution buffer. The enriched chromatin was analyzed by qPCR using b-actin primers, as described (Kaneko et al., 2007): Promoter (Prom): CTC AAT CTC GCT CTC GCT CT and CTC GAG CCA TAA AAG GCA AC; Region B: CAA CTG GGA CGA CAT GGA GAA A and GAG TCC TAC GG AAA ACG GCA GA; Poly (A): TGT ACA CTG ACT TGA GAC CAG T and AAG CAG GAA CAG AGA CCT GAC C; Region D: TAG GCT TAG GAG AGG CCG CAA T and GTC CAG GAG CCT GGG TAT CTC C. For ChIP Pol II results, the relative signal was normalized to the IgG, and for ChIP XRN2 results, the signal was normalized to the Pol II.

### *In vitro* methylation Assay

GST-tagged DDX5 fragments and mutants were purified from bacteria. Ten µg of each GST-tagged construct was incubated with 2 µl of (methyl-^3^H) S-adenosyl-L-methionine solution (15Ci/mmol stock solution, 0.55 µM final concentration, Perkin-Elmer) and 2 µg of PRMT5:MEP50 active complex (Sigma Aldrich) in methylation buffer (HEPES pH 7.4 50mM, NP40 0,01%, DTT 1 mM, PMSF 1 mM) for 2h at 37°C. Samples were separated by SDS-page and stained with Coomassie Blue. After de-staining, the gel was then incubated for 1h in EN^3^HANCE (Perkin Elmer), according to manufacturer's instructions and the reaction was visualized by fluorography.

### FACS-Based Cell Survival Assay

For FACS-based cell survival assay, a U2OS cell line with stably transfected GFP (U2OS-GFP) was generated. The U2OS and U2OS-GFP cells were transfected with control and target siRNAs respectively, or reversely the U2OS cells transfected with target siRNAs and U2OS-GFP cells with control siRNA. The cells were trypsinized and mixed with an approximately 1:1 ratio two days after transfection. The cells were then co-plated and treated with DNA damage agents or left untreated. After seven to ten days of recovery, the cells were subjected to FACS analysis to determine the ratio of GFP+/GFP-(green/non-green) cells which reflects the relative survival of the two cell populations.

### *In vitro* Unwinding Assay

DDX5 and XRN2 were tagged at the N-terminus with GST and at the C-terminus with His_10_, expressed in bacteria and purified as described for PALB2 (Buisson, Niraj et al., 2014). R-loop and D-loop substrates were generated by annealing purified oligonucleotides: DNA strand 1: 5’GGGTGAACCTGCAGGTGGGCGGCTGCTCATCGTAGGTTAGTTGGTAGAATTCGGCAGCGTC-3’ DNA strand 2: 5’-GACGCTGCCGAATTCTACCAGTGCCTTGCTAGGACATCTTTGCCCACCTGCAGGTTCACCC-3’ With either RNA strand: 5’-AAAGAUGUCCUAGCAAGGCAC-3’ or DNA strand: 5’-AAAGATGTCCTAGCAAGGCAC-3’. Unwinding assays were performed in MOPS buffer (25 mM MOPS (morpholinepropanesulfonic acid) pH 7.0, 60 mM KCl, 0.2% Tween-20, 2 mM DTT, 5 mM ATP, 5 mM MgCl_2_). DDX5 and labelled R-loop or D-loop (100 nM) substrates were incubated in MOPS buffer for 20 minutes at 37°C, followed by deproteinization in one-fifth volume of stop buffer (20 mM Tris-Cl pH 7.5 and 2 mg/mL proteinase K) for 20 min at 37°C. Reactions were loaded on an 8 % acrylamide gel, electrophoresed at 150V for 120 min, dried onto filter paper and autoradiographed.

### R-loop RNase Assays

XRN2 RNase assays were performed in Tris/MOPS buffer (12.5 mM MOPS (morpholinepropanesulfonic acid) pH 7.0, 25 mM TRIS-HCl pH 7.9, 30 mM KCl, 50 mM NaCl, 0.1% Tween-20, 1.5 mM DTT, 5 mM ATP, 10 mM MgCl_2_). DDX5 and labelled R-loop (100 nM) substrates were incubated in TRIS/MOPS buffer for 20 min at 37°C, XRN2 was added for 20 min at 37°C. Reactions were deproteinized in one-fifth volume of stop buffer (20 mM Tris-Cl pH 7.5 and 2 mg/mL proteinase K) for 20 min at 37°C. Reactions were loaded on an 8 % acrylamide gel, electrophoresed at 150V for 120 min, dried onto filter paper and autoradiographed.

### Clonogenic cell survival assay

For clonogenic assay, 200-1000 cells/10-cm dish were seeded and the cells were treated with hydroxyurea (HU) for 20h. Ten to Fourteen days after the treatment, Cells were fixed with 4% paraformaldehyde and stained with 0.05% crystal violet (Sigma-Aldrich) and colonies were counted.

## Author Contributions

Z.Y., S.Y.M, J.Y.M and S.R. designed the research; Z.Y., S.Y.M, Y.C, M.K. and F.F.B. performed the experiments; Z.Y., S.Y.M, M.K. J.Y.M and S.R analyzed the data; Z.Y., J.Y.M and S.R. wrote the paper.

## Competing Interests

The authors declare no competing interests.

## References

Adamson B, Smogorzewska A, Sigoillot FD, King RW, Elledge SJ (2012) A genome-wide homologous recombination screen identifies the RNA-binding protein RBMX as a component of the DNA-damage response. Nat Cell Biol 19: 318–328

Aguilera A, Gomez-Gonzalez B (2017) DNA-RNA hybrids: The risks of DNA breakage during transcription. Nat Struct Mol Biol 24: 439–443

Aymard F, Aguirrebengoa M, Guillou E, Javierre BM, Bugler B, Arnould C, Rocher V, Iacovoni JS, Biernacka A, Skrzypczak M, Ginalski K, Rowicka M, Fraser P, Legube G (2017) Genome-wide mapping of long-range contacts unveils clustering of DNA double-strand breaks at damaged active genes. Nat Struct Mol Biol 24: 353–361

Bedford MT, Clarke SG (2009) Protein arginine methylation in mammals: who, what, and why. Mol Cell 33: 1–13

Bezzi M, Teo SX, Muller J, Mok WC, Sahu SK, Vardy LA, Bonday ZQ, Guccione E (2013) Regulation of constitutive and alternative splicing by PRMT5 reveals a role for Mdm4 pre-mRNA in sensing defects in the spliceosomal machinery. Genes Dev 27: 1903–1916

Bhatia V, Barroso SI, García-Rubio ML, Tumini E, Herrera-Moyano E, Aguilera A (2014) BRCA2 prevents R-loop accumulation and associates with TREX-2 mRNA export factor PCID2. Nature 511: 362–365

Bhatia V, Herrera-Moyano E, Aguilera A, Gomez Gonzaleze B (2017) The role of replication-associated repair factors on R-loops. Genes 8: 171

Blanc RS, Richard S (2017) Arginine methylation: the coming of Age. Molecular Cell 65: 8–24

Branscombe TL, Frankel A, Lee JH, Cook JR, Yang Z, Pestka S, Clarke S (2001) PRMT5 (Janus kinase-binding protein 1) catalyzes the formation of symmetric dimethylarginine residues in proteins. J Biol Chem 276: 32971–32976

Braun CJ, Stanciu M, Boutz PL, Patterson JC, Calligaris D, Higuchi F, Neupane R, Fenoglio S, Cahill DP, Wakimoto H, Agar NYR, Yaffe MB, Sharp PA, Hemann MT, Lees JA (2017) Coordinated Splicing of Regulatory Detained Introns within Oncogenic Transcripts Creates an Exploitable Vulnerability in Malignant Glioma. Cancer cell 32: 411–426

Buisson R, Niraj J, Pauty J, Maity R, Zhao W, Coulombe Y, Sung P, Masson JY (2014) Breast cancer proteins PALB2 and BRCA2 stimulate polymerase η in recombination-associated DNA synthesis at blocked replication forks. Cell Rep 6: 553–564

Chan-Penebre E, Kuplast KG, Majer CR, Boriack-Sjodin PA, Wigle TJ, Johnston LD, et al. (2015) A selective inhibitor of PRMT5 with in vivo and in vitro potency in MCL models. Nature Chemical Biology 11: 432–437

Chedin F (2016) Nascent connections: R-loops and chromatin patterning. Trends Genet 32: 828–838

Cho E-C, Zheng S, Munro S, Liu G, Carr SM, Moehlenbrink J, Lu Y-C, Stimson L, Khan O, Konietzny R, McGouran J, Coutts AS, Kessler B, Kerr DJ, Thangue NBL (2012) Arginine methylation controls growth regulation by E2F-1. EMBO J 31: 1785–1797

Choi YJ, Lee SG (2012) The DEAD-box RNA helicase DDX3 interacts with DDX5, co-localizes with it in the cytoplasm during the G2/M phase of the cycle, and affects its shuttling during mRNP export. Journal of Cellular Biochemistry 113: 985–996

Clarke TL, Sanchez-Bailon MP, Chiang K, Reynolds JJ, Herrero-Ruiz J, Bandeiras TM, Matias PM, Maslen SL, Skehel JM, Stewart GS, Davies CC (2017) PRMT5-Dependent Methylation of the TIP60 Coactivator RUVBL1 Is a Key Regulator of Homologous Recombination. Mol Cell 65: 900–916

Cloutier SC, Wang S, Ma WK, Al Husini N, Dhoondia Z, Ansari A, Pascuzzi PE, Tran EJ (2016) Regulated Formation of lncRNA-DNA Hybrids Enables Faster Transcriptional Induction and Environmental Adaptation. Mol Cell 62: 148

Cohen S, Puget N, Lin YL, Clouaire T, Aguirrebengoa M, Rocher V, Pasero P, Canitrot Y, Legube G (2018) Senataxin resolves RNA:DNA hybrids forming at DNA double-strand breaks to prevent translocations. Nat Commun 9: 533

Cristini A, Groh M, Kristiansen MS, Gromak N (2018) RNA/DNA Hybrid Interactome Identifies DXH9 as a Molecular Player in Transcriptional Termination and R-Loop-Associated DNA Damage. Cell Rep 23: 1891–1905

Francia S, Michelini F, Saxena A, Tang D, de Hoon M, Anelli V, Mione M, Carninci P, d'Adda di Fagagna F (2012) Site-specific DICER and DROSHA RNA products control the DNA-damage response. Nature 488: 231–235

Friesen WJ, Massenet S, Paushkin S, Wyce A, Dreyfuss G (2001) SMN, the product of the spinal muscular atrophy gene, binds preferentially to dimethylarginine-containing protein targets. Mol Cell 7: 1111–1117

Fuller-Pace FV (2013) The DEAD box proteins DDX5 (p68) and DDX17 (p72): multi-tasking transcriptional regulators. Biochimica et biophysica acta 1829: 756–763

García-Rubio ML, Pérez-Calero C, Barroso SI, Tumini E, Herrera-Moyano E, Rosado IV, Aguilera A (2015) The Fanconi Anemia Pathway Protects Genome Integrity from R-loops. PLoS Genet 11: e1005674

Ginno PA, Lott PL, Christensen HC, Korf I, Chedin F (2012) R-loop formation is a distinctive characteristic of unmethylated human CpG island promoters. Mol Cell 45: 814–825

Hamard PJ, Santiago GE, Liu F, Karl DL, Martinez C, Man N, Mookhtiar AK, Duffort S, Greenblatt S, Verdun RE, Nimer SD (2018) PRMT5 Regulates DNA Repair by Controlling the Alternative Splicing of Histone-Modifying Enzymes. Cell Rep 24: 2643–2657

Hamperl S, Bocek MJ, Saldivar JC, Swigut T, Cimprich KA (2017) Transcription-Replication Conflict Orientation Modulates R-Loop Levels and Activates Distinct DNA Damage Responses. Cell 170: 774–786

Hamperl S, Cimprich KA (2016) Conflict resolution in the genome: How transcription and replication make it work. Cell 167: 1455–1467

Hatchi E, Skourti-Stathaki K, Ventz S P, L., Yen A, Kamieniarz-Gdula K, Dimitrov S, Pathania S, McKinney KM, Eaton ML, Kellis M, Hill SJ, Parmigiani G, Proudfoot NL, Livingston DM (2015) BRCA1 recruitment to transcriptional pause sites is required for R-loop-driven DNA damage repair. Mol Cell 57: 636–647

Hirling H, Scheffner M, Restle T, Stahl H (1989) RNA helicase activity associated with the human p68 protein. Nature 339: 562–564

Hodroj D, Recolin B, Serhal K, Martinez S, Tsanov N, Abou Merhi R, Maiorano D (2017) An ATR-dependent function for the Ddx19 RNA helicase in nuclear R-loop metabolism. EMBO J 36: 1182–1198

Huertas P, Aguilera A (2003) Cotranscriptionally formed DNA:RNA hybrids mediate transcription elongation impairment and transcription-associated recombination. Mol Cell 12: 711–721

Kaneko S, Rozenblatt-Rosen O, Meyerson M, Manley JL (2007) The multifunctional protein p54nrb/PSF recruits the exonuclease XRN2 to facilitate pre-mRNA 3' processing and transcription termination. Genes Dev 21: 1779–1789

Kaushik S, Liu F, Veazey KJ, Gao G, Das P, Neves LF, Lin K, Zhong Y, Lu Y, Giuliani V, Bedford MT, Nimer SD, Santos MA (2018) Genetic deletion or small-molecule inhibition of the arginine methyltransferase PRMT5 exhibit anti-tumoral activity in mouse models of MLL-rearranged AML. Leukemia 32: 499–509

Kim M, Krogan NJ, Vasiljeva L, Rando OJ, Nedea E, Greenblatt JF, Buratowski S (2004) The yeast Rat1 exonuclease promotes transcription termination by RNA polymerase II. Nature 432: 517–522

Koh CM, Bezzi M, Low DH, Ang WX, Teo SX, Gay FP, Al-Haddawi M, Tan SY, Osato M, Sabò A, Amati B, Wee K, Guccione E (2015) MYC regulates the core pre-mRNA splicing machinery as an essential step in lymphomagenesis. Nature 523: 96–100

Li L, Germain DR, Poon HY, Hildebrandt MR, Monckton EA, McDonald D, Hendzel MJ, Godbout R (2016) DEAD Box 1 Facilitates Removal of RNA and Homologous Recombination at DNA Double-Strand Breaks. Mol Cell Biol 36: 2794–2810

Li L, Monckton EA, Godbout R (2008) A role for DEAD box 1 at DNA double-strand breaks. Mol Cell Biol 28: 6413–6425

Li X, Manley JL (2005) Inactivation of the SR protein splicing factor ASF/SF2 results in genomic instability. Cell 122: 365–378

Li Y, Chitnis N, Nakagawa H, Kita Y, Natsugoe S, Yang Y, Li Z, Wasik M, Klein-Szanto AJ, Rustgi AK, Diehl JA (2015) PRMT5 is required for lymphomagenesis triggered by multiple oncogenic drivers. Cancer Discov 5: 288–303

Morales JC, Richard P, Patidar PL, Motea EA, Dang TT, Manley JL, Boothman DA (2016) XRN2 Links Transcription Termination to DNA Damage and Replication Stress. PLoS Genet 12: e1006107

Nicol SM, Bray SE, Black HD, Lorimore SA, Wright EG, Lane DP, Meek DW, Coates PJ, Fuller-Pace F (2013) The RNA helicase p68 (DDX5) is selectively required for the induction of p53-dependent p21 expression and cell-cycle arrest after DNA damage. Oncogene 32: 3461–3469

Ogilvie VC, Wilson BJ, Nicol SM, Morrice NA, Saunders LR, Barber GN, Fuller-Pace FV (2003) The highly related DEAD box RNA helicases p68 and p72 exist as heterodimers in cells. Nucleic Acids Research 31: 1470–1480

Pal S, Vishwanath SN, Erdjument-Bromage H, Tempst P, Sif S (2004) Human SWI/SNF-associated PRMT5 methylates histone H3 arginine 8 and negatively regulates expression of ST7 and NM23 tumor suppressor genes. Mol Cell Biol 24: 9630–9645

Paulsen RD, Soni DV, Wollman R, Hahn AT, Yee MC, Guan A, Hesley JA, Miller SC, Cromwell EF, Solow-Cordero DE, al. e (2009) A genome-wide siRNA screen reveals diverse cellular processes and pathways that mediate genome stability. Mol Cell 35: 228–239

Ribeiro de Almeida C, Dhir S, Dhir A, Moghaddam AE, Sattentau Q, Meinhart A, Proudfoot N (2018) RNA Helicase DDX1 Converts RNA G-Quadruplex Structures into R-Loops to Promote IgH Class Switch Recombination. Mol Cell 70: 650–662

Rossler OG, Straka A, Stahl H (2001) Rearrangement of structured RNA via branch migration structures catalysed by the highly related DEAD-box proteins p68 and p72. Nucleic Acids Research 29: 2088–2096

Schwab RA, Nieminuszczy J, Shah F, Langton J, Lopez Martinez D, Liang CC, Cohn MA, Gibbons RJ, Deans AJ, Niedzwiedz W (2015) The Fanconi Anemia Pathway Maintains Genome Stability by Coordinating Replication and Transcription. Mol Cell 60: 351–361

Shanbhag NM, Rafalska-Metcalf IU, Balane-Bolivar C, Janicki SM, Greenberg RA (2010) An ATM-Dependent Transcriptional Silencing Program is Transmitted Through Chromatin in Cis to DNA Double Strand Breaks. Cell 141: 970–981

Silva S, Camino LP, Aguilera A (2018) Human mitochondrial degradosome prevents harmful mitochondrial R loops and mitochondrial genome instability. Proceedings of the National Academy of Sciences of the United States of America doi: 10.1073/pnas.1807258115.

Skourti-Stathaki K, Proudfoot NJ (2014) A double-edged sword: R loops as threats to genome integrity and powerful regulators of gene expression. Genes Dev 28: 1384–1396

Skourti-Stathaki K, Proudfoot NJ, Gromak N (2011) Human senataxin resolves RNA/DNA hybrids formed at transcriptional pause sites to promote Xrn2-dependent termination. Mol Cell 42: 794–805

Sollier J, Stork CT, García-Rubio ML, Paulsen RD, Aguilera A, Cimprich KA (2014) Transcription-coupled nucleotide excision repair factors promote R-loop-induced genome instability. Mol Cell 56: 777–785

Song C, Hotz-Wagenblatt A, Voit R, Grummt I (2017) SIRT7 and the DEAD-box helicase DDX21 cooperate to resolve genomic R loops and safeguard genome stability. Genes Dev 31: 1370–1381

Sridhara SC, Carvalho S, Grosso AR, Gallego-Paez LM, Carmo-Fonseca M, de Almeida SF (2017) Transcription Dynamics Prevent RNA-Mediated Genomic Instability through SRPK2-Dependent DDX23 Phosphorylation. Cell Rep 18: 334–343

Stirling PC, Chan YA, Minaker SW, Aristizabal MJ, Barrett I, Sipahimalani P, Kobor MS, Hieter P (2012) R-loop-mediated genome instability in mRNA cleavage and polyadenylation mutants. Genes Dev 26: 163–175

Thandapani P, O'Connor TR, Bailey TL, Richard S (2013) Defining the RGG/RG motif. Mol Cell 50: 613–623

Tuduri S, Crabbé L, Conti C, Tourrière H, Holtgreve-Grez H, Jauch A, Pantesco V, De Vos J, Thomas A, Theillet C, Pommier Y, Tazi J, Coquelle A, Pasero P (2009) Topoisomerase I suppresses genomic instability by preventing interference between replication and transcription. Nat Cell Biol 11: 1315–1324

Wahba L, Amon JD, Koshland D, Vuica-Ross M (2011) RNase H and multiple RNA biogenesis factors cooperate to prevent RNA:DNA hybrids from generating genome instability. Mol Cell 44: 978–988

Wahba L, Gore SK, Koshland D (2013) The homologous recombination machinery modulates the formation of RNA-DNA hybrids and associated chromosome instability. Elife 2: e00505

Wang IX, Grunseich C, Fox J, Burdick J, Zhu Z, Ravazian N, Hafner M, Cheung V (2018) Human proteins that interact with RNA/DNA hybrids. Genome Res 28: 1405–1414

West S, Gromak N, Proudfoot NJ (2004) Human 5' --> 3' exonuclease Xrn2 promotes transcription termination at co-transcriptional cleavage sites. Nature 432: 522–525

Wilson BJ, Giguere V (2007) Identification of novel pathway partners of p68 and p72 RNA helicases through Oncomine meta-analysis. BMC Genomics 8: 419

Xing Z, Wang S, Tran EJ (2017) Characterization of the mammalian DEAD-box protein DDX5 reveals functional conservation with S. cerevisiae ortholog Dbp2 in transcriptional control and glucose metabolism. RNA (New York, NY) 23: 1125–1138

Yang Y, McBride KM, Hensley S, Lu Y, Chedin F, Bedford MT (2014) Arginine methylation facilitates the recruitment of TOP3B to chromatin to prevent R loop accumulation. Mol Cell 53: 484–497

Zhao DY, Gish G, Braunschweig U, Li Y, Ni Z, Schmitges FW, Zhong G, Liu K, Li W, Moffat J, Vedadi M, Min J, Pawson TJ, Blencowe BJ, Greenblatt JF (2016) SMN and symmetric arginine dimethylation of RNA polymerase II C-terminal domain control termination. Nature 529: 48–53

Zhou J, Xu F, Jin B, Cui L, Wang Y, Du X, Li J, Li P, Ren R, Pan J (2016) Targeting methyltransferase PRMT5 eliminates leukemia stem cells in chronic myelogenous leukemia. Journal of Clinical Investigation 126: 3961–3980

